# Seed microbiota revealed by a large-scale meta-analysis including 50 plant species

**DOI:** 10.1101/2021.06.08.447541

**Authors:** Marie Simonin, Martial Briand, Guillaume Chesneau, Aude Rochefort, Coralie Marais, Alain Sarniguet, Matthieu Barret

**Affiliations:** Univ Angers, Institut Agro, INRAE, IRHS, SFR QUASAV, F-49000 Angers, France

**Keywords:** Plant microbiome, Diversity, Data synthesis, Metabarcoding, Fungal community

## Abstract

Seed microbiota constitutes a primary inoculum for plants that is gaining attention due to its role for plant health and productivity. Here, we performed a meta-analysis on 63 seed microbiota studies covering 50 plant species to synthesize knowledge on the diversity of this habitat. Seed microbiota are diverse and extremely variable, with taxa richness varying from one to thousands of taxa. Hence, seed microbiota presents a variable (i.e flexible) microbial fraction but we also identified a stable (i.e. core) fraction across samples. Around 30 bacterial and fungal taxa are present in most plant species and in samples from all over the world. Core taxa, such as *Pantoea agglomerans, Pseudomonas viridiflava, P. fluorescens, Cladosporium perangustum* and *Alternaria sp*., are dominant seed taxa. The characterization of the core and flexible seed microbiota provided here will help uncover seed microbiota roles for plant health and design effective microbiome engineering.

## INTRODUCTION

Seeds are key components of plant fitness and are central to the sustainability of the agri-food system. Seed consumption is the foundation of human food security with wheat, rice and maize seeds providing 42.5 % of the world’s food calorie supply^1^. Moreover, the transition of seed to seedling represents a major bottleneck for both plant fitness and seed microbiota with large implications in agricultural systems and for the maintenance of plant biodiversity in natural ecosystems^2–4^. Both seed quality for food consumption and seed vigour in agricultural settings can be influenced by the microorganisms living inside and on the surface of seeds (i.e. the seed microbiota)^5,6^. Knowledge regarding seed microbiota has been long lagging behind that of other plant compartments, like the rhizosphere, phyllosphere and endosphere^7^. However, recently a renewed attention to reproductive tissues (seeds, flowers) and plant early life stages have emerged to better understand the dynamic and assembly processes of plant microbiota ^8–10^.

Seed microbiota constitutes a primary microbial inoculum for the plant microbiota with potential long-term impacts on plant fitness. Seed-associated microorganisms can be acquired either horizontally from various environments (e.g. air, water, insects, seed processing) or vertically from the mother plant and transmitted across multiple generations ^11–13^. The prevalence of pathogens on seeds has been extensively studied but the occurrence and role of other commensal or mutualistic microorganisms constituting the majority of seed microbiota are mostly unknown ^14–16^. Currently, limited attempts have been made to characterize the seed core microbiota of a specific plant species or shared across multiple plant species at a large scale ^17,18^. Here, the definition of the core microbiota used, corresponds to the “common core” (*sensu* Risely 2020^19^) that represents the component of the microbiota that is found across a considerable proportion of hosts. The identification of the core and flexible microbiota of a plant habitat can help identify microbial taxa and functions that may be particularly important for host fitness^20^. This identification can be achieved by large-scale data synthesis efforts (e.g. meta-analysis) but such efforts remain to be done for seed microbiota.

A global analysis of seed microbial diversity appears even more timely as seeds appear as a key vector of solutions to promote sustainable agriculture. Seeds can play a double role: i) as a source of innovations with seed-borne microorganisms representing key biotechnological resources^21^; and ii) as carriers of microbial biostimulants or biocontrol solutions that greatly reduces the surface and volume of treatment applied to fields, thus decreasing application costs and potential negative impacts on the environment ^22–24^. A synthesized knowledge on seed microbiota will accelerate discoveries and will help future practices promoting the presence of important seed microorganisms for plant health and productivity.

In this context, we performed a meta-analysis on available seed microbiota studies to synthesize the current knowledge on the diversity of this habitat and to constitute an open database for the research community. This data synthesis effort enabled us to address the following questions:

1. How diverse is the seed microbiota?
2. Which taxonomic groups compose the seed microbiota?
3. Can we find the evidence for a seed core microbiota shared across plant species?
4. Do we detect specific patterns in seed microbiota composition and diversity by plant species?

These questions were addressed through a meta-analysis gathering a total of 63 seed microbiota studies yielding 3,190 seed samples from 50 plant species collected in 28 countries. This study shows that the overall seed microbiota composition is highly variable from one seed sample to another but most seeds share a core microbiota composed of few dominant bacterial and fungal taxa.

## RESULTS

### Creation of the Seed Microbiota Database to perform the meta-analysis

To synthesize knowledge on the diversity and composition of seed microbiota, we identified, re-processed and re-analyzed raw data from published microbiome studies and also unpublished data from our laboratory. We initially identified 100 seed microbiota studies (59 for 16S rRNA gene; 14 for *gyrB* and 27 for fungal Internal Transcribed Spacer (ITS) regions) that used amplicon sequencing (i.e metabarcoding) to characterize community structure. To enable the comparison of different studies and the use of a common bioinformatic pipeline, only studies using the Illumina sequencing technology were considered (n=90). A total of 63 studies were finally included in the publicly available database that we called Seed Microbiota Database, based on the (i) availability of fastq files and associated metadata, (ii) sequencing quality and (iii) primer sets employed (**Fig. 1**). The Seed Microbiota Database is available on the Data INRAE portal (https://doi.org/10.15454/2ANNJM). Detailed information on the 63 studies and references can be found in Supplementary Data 1 and **Fig. 1C**^4,11,13,17,25–47^. Each study was independently re-processed with a standardized bioinformatic pipeline (https://github.com/marie-simonin/Seed-Microbiota-Database) using Qiime2 and DADA2 before being merged with other studies targeting the same marker gene and region to form a final “dataset”. A total of 5 final datasets were included in the meta-analysis, which corresponds to the most common molecular markers (16S rRNA-V4, 16S rRNA-V5-V6, *gyrB*, ITS1 and ITS2) used to characterize seed microbiomes (**Fig. 1B**). Detailed information on the description of the samples (e.g. origin, plant species, seed preparation, seed fraction, seed number) for the different molecular markers employed can be found in supplementary materials (**Fig. S1, S2 and S3**).

**Figure 1:**
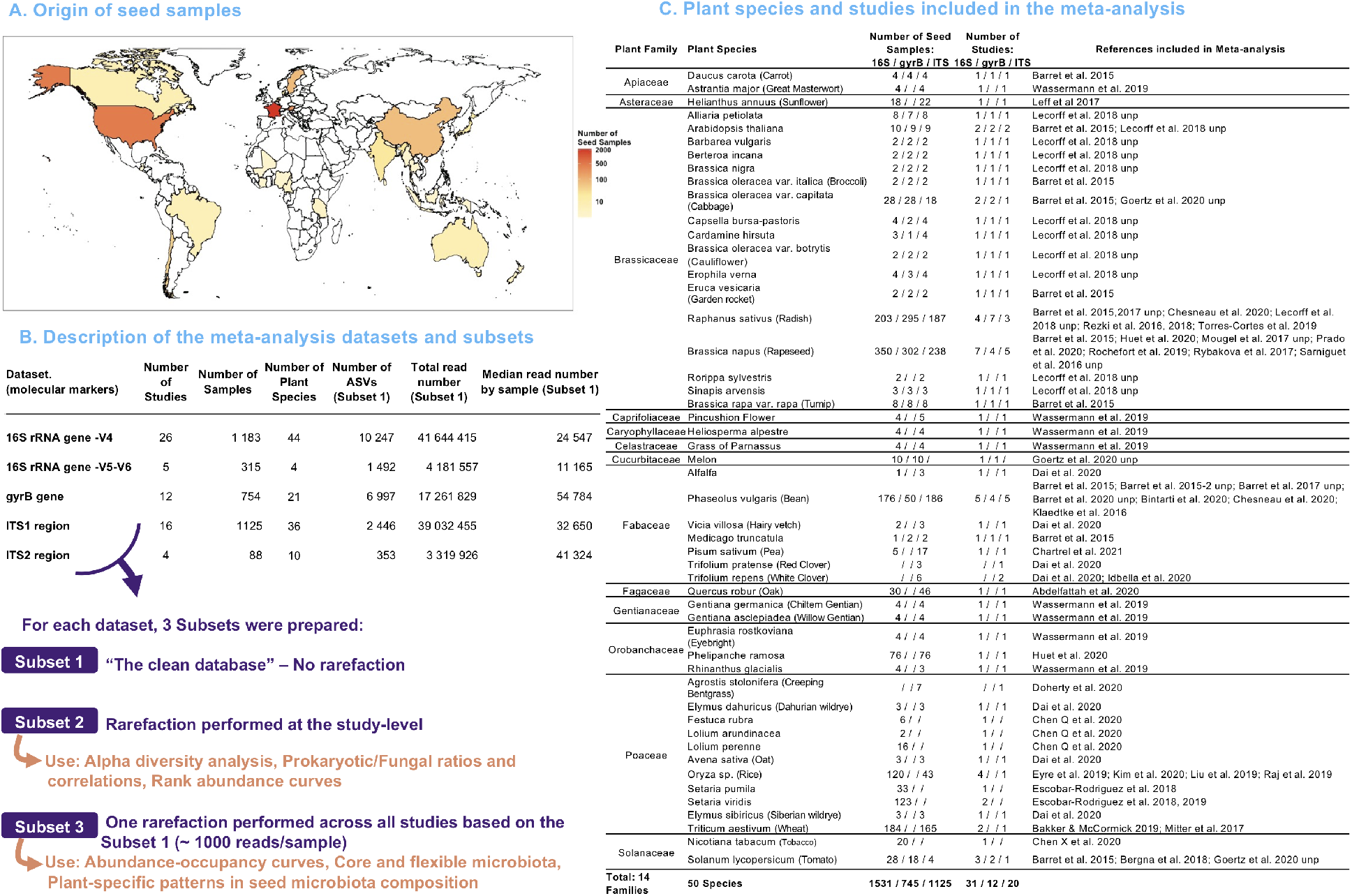
Overview of the seed microbiota meta-analysis database. A) Map presenting the countries of origin of seed samples (n=28 countries). The countries are colored based on the number of seed samples that originated from this area. B) Table providing general information on the 5 datasets (16S rRNA-V4, 16S rRNA-V5-V6, *gyrB*, ITS1 and ITS2) of the meta-analysis. Each of these 5 datasets were prepared in three different subsets adapted to different types of downstream analyses. C) Table presenting for each plant species included in the meta-analysis (n=50 plant species): the number of samples and studies for the three marker genes and the references of the studies. “unp”: unpublished dataset originating from our research group.

In the end, this meta-analysis was conducted on a total of 3,190 samples from 50 plant species collected in 28 countries. The community profiles of these samples were estimated *via* three bacterial markers (*gyrB*, V4 and V5-V6 regions of 16S rRNA gene) and two fungal markers (ITS1 and ITS2) with 105 millions of paired-end reads and thousands of Amplicon Sequence Variants (ASVs) identified (**Fig. 1**). The 16S-V4, *gyrB* and ITS1 datasets presented the highest number of samples and plant diversity (**Fig. 1B**) and thus have been primarily used for community profiling of the seed microbiota in this meta-analysis. Each dataset was prepared in three different subsets adapted to different types of downstream analyses (Subset 1: no rarefaction; Subset 2: rarefaction at the study-level; Subset 3: rarefaction across all studies, **Fig. 1B** and **Table S1**).

The seed samples included in the meta-analysis cover an important diversity of plant families (n=14). Still, it should be noted that 73% of all samples came from four species from Brassicaceae (*Raphanus sativus* (radish), *Brassica napus* (rapeseed)), Fabaceae (*Phaseolus vulgaris* (bean)) and Poaceae (*Triticum aestivum* (wheat)) (**Fig. 1C**, **Fig. S1, S2, S3**). Most of the other plant species investigated were covered by less than 40 seed samples (n=37), with the exception of *Oryza sp*. (rice), *Setaria viridis, Phelipanche ramosa, Solanum lycopersicum (tomato), Brassica oleracea var. capitata (cabbage)* and *Helianthus annuus* (sunflower) (**Fig. S1, S2, S3**). Seed samples were harvested across all continents with a significant number of samples from France (n=2159), the USA (n=457), Austria (n=261), Sweden (n=79) and China (n=71), and were analyzed by 15 research institutes/universities. Different procedures were employed for characterizing seed microbiota structure including raw seed samples (i.e. total fraction), surface-disinfected, surface-washed and dissected seeds. Additionally, seeds were prepared following different approaches before DNA extractions (e.g soaking, grinding, washing, **Fig. S1, S2, S3**). All studies except one (Abdelfattah et al. 2021, *Quercus robur L*. (oak)) characterized seed microbiota structure on seed lots (> 3 seeds) because the microbial DNA present on a single seed is often not enough to perform metabarcoding^48^. Hence, seed microbiota were characterized on samples with a variable number of seeds depending on the study, but the majority of samples were composed of 1000 seeds providing a view of the seed “metacommunity” of a given site.

### Part 1: Patterns across all plant species

#### a) Highly variable seed microbiota diversity

The alpha diversity patterns of seed microbiota were explored using observed ASV richness, Pielou’s evenness and Shannon diversity index. A large variability in ASV richness per seed sample was observed within and across plant species (16S rRNA gene markers: 1 to 2224 ASVs, *gyrB*: 3 to 718 ASVs, ITS markers: 1 to 423 ASVs, **Fig. 2**). The median ASV richness across all samples were 48 ± 8 for the 16S rRNA gene markers, 63 ± 8.6 for *gyrB* and 52 ± 2.7 for ITS markers indicating similar levels of prokaryotic (Archaea + Bacteria) and fungal richness in seed microbiota. When considering only the plant species that had a high number of samples and several independent studies, we observed that tomato had one of the highest prokaryotic richness (16S rRNA gene, median: 266 ASVs), rapeseed presented intermediate levels (median: 57 ASVs), while rice (33 ASVs), radish (23 ASVs) and bean seeds (23 ASVs) were on the lower end. For fungal richness, rapeseed (median: 67 ASVs) and bean (67 ASVs) were on the higher end, while radish seeds presented intermediate levels (51 ASVs). A similar variability between seed samples was observed in Pielou’s evenness values (**Fig. S4**) and Shannon index (**Fig. S5**). Between samples from the same study, seed bacterial and fungal communities can range from being extremely uneven (<0.25) to highly even (>0.75) for several plant species, such as bean, radish, rapeseed and wheat (**Fig. S4**).

**Figure 2:**
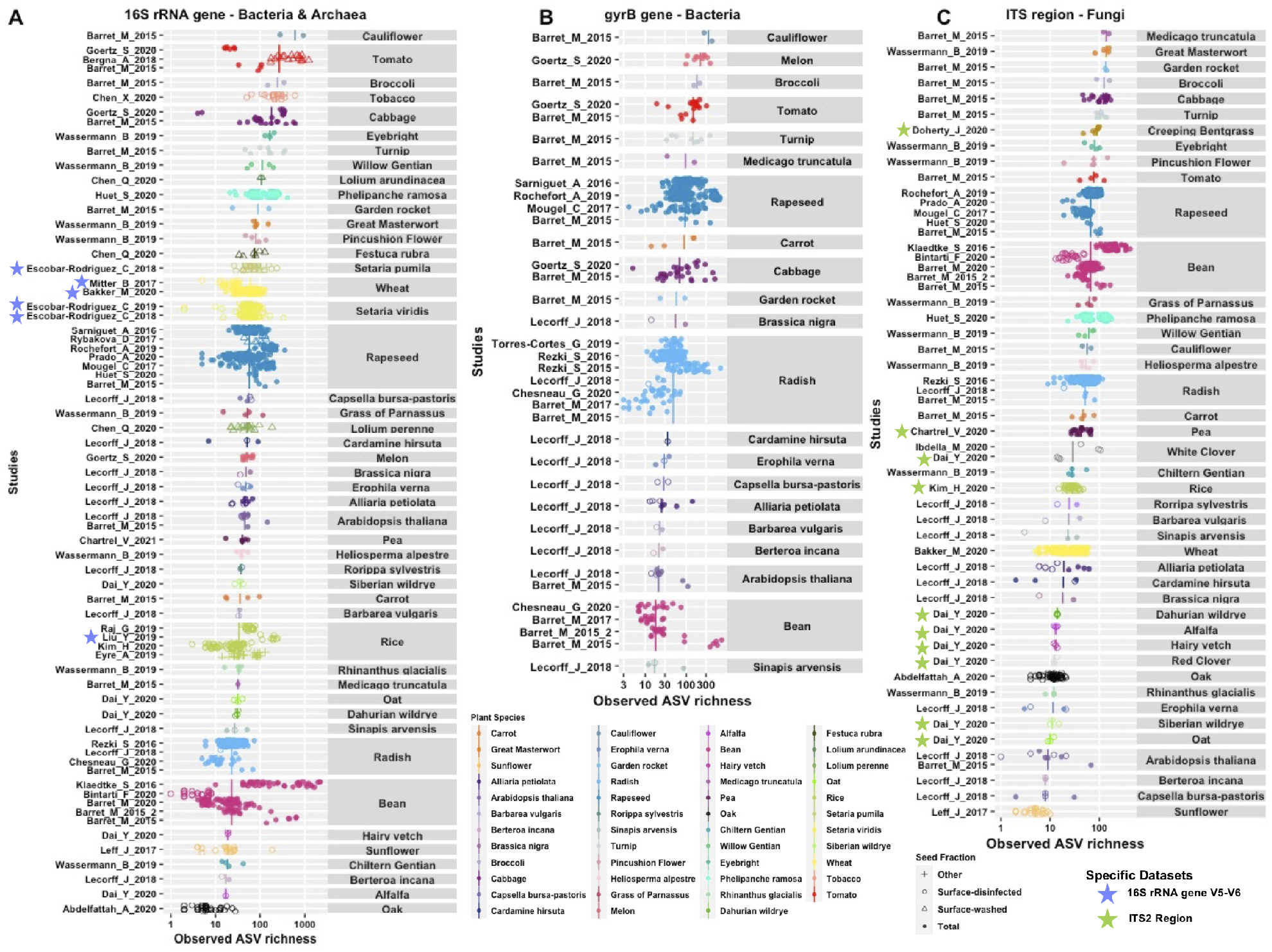
Seed microbiota diversity patterns within and across plant species. Observed ASV richness of all the seed samples included in the meta-analysis for the 16S rRNA gene markers (panel A, n=1531, V4 and V5-V6 datasets combined), *gyrB* (panel B, n=754) and ITS markers (panel C, n=1125, ITS1 and ITS2 datasets combined). The few 16S V5-V6 studies are highlighted with a blue star and the ITS2 studies with a green star. Each point represents a seed sample that can be composed of multiple seeds (up to a thousand seeds). The shape of the points represents the seed fraction considered (Total: no pre-treatment of the seeds, Surface-washed: seeds were rinsed with sterile water, Surface-disinfected: seeds were surface sterilized with chemical products, Other: seeds were dissected or received specific treatments). For each plant species, the vertical line corresponds to the median value of the samples. Note that the *x* axis is presented on a log scale and that the range differs on each panel. On each panel, species are ordered based on the median richness value of the plant species (lowest richness at the bottom and highest at the top).

We also considered the influence of the seed fraction studied, as many studies performed a surface-disinfection of the seeds to enrich the endophytic fraction of the microbiota. The median ASV richness of surface-disinfected seeds was only slightly lower (40 ± 6, n=415) than non-disinfected seeds (Total fraction, 50 ± 10, n=1039) for the 16S rRNA gene samples but the loss of ASVs with surface-disinfection was more pronounced for the ITS samples (16 ± 2, n=193 vs 47 ± 3, n=930, **Fig. S6**). Regarding the seed preparation technique used before DNA extraction, similar levels of prokaryotic ASV richness were observed between the different techniques except for the fungal ASV richness that was higher with seed soaking (61 ± 3, n=802) compared to grinding (21 ± 2, n=322, **Fig. S7**). However, this latter result could be mainly driven by the fact that different plant species were studied with different techniques. We also explored if the variability observed in seed microbiota diversity was associated with the number of seeds included in the seed sample (from 1 to 1000 seeds). We found that ASV richness generally increased with the number of seeds in the sample for both prokaryotic and fungal communities, but a high variability still existed for each given seed sample size (e.g. 1000 seeds, **Fig. S8).**

Altogether, these results show that seed microbiota are diverse with a median of 100 ASVs ± 11 by sample (sum of prokaryotic and fungal diversity) and are also extremely variable from one sample to another even within the same study or for a given plant species. These analyses also suggest that seed surface-disinfection and preparation have more impacts on the fungal community than the prokaryotic community.

Next, we explored the relationships between the prokaryotic and fungal richness to determine the group of dominance and their diversity correlation. This analysis was performed on samples for which both 16S rRNA gene and ITS metabarcoding were performed (n=884). The prokaryotic and fungal ASV richness were weakly positively correlated (R^2^=0.11, P<0.001, **Fig. 3A**). Seed microbiota presented a variable proportion of bacterial and fungal taxa but interestingly communities were not skewed towards one particular microbial group (**Fig. 3A**). Values were distributed close to the 1 to 1 line (similar diversity in both groups) or on both sides of the lines for most species and plant families (**Fig. 3A**, **3B**). These results indicate that seed microbiota can be either dominated by fungal or prokaryotic taxa in terms of diversity. Only Poaceae (3 studies) and Orobanchaceae (2 studies) seeds appeared to present a higher proportion of prokaryotic richness (**Fig. 3A, 3B**). Still, large differences in the proportion of prokaryotic and fungal richness were observed between and among plant species (**Fig. 3B**). For instance, Brassicaceae species ranged from being dominated by fungal taxa (e.g. radish) to being highly dominated by prokaryotic taxa (e.g. cauliflower, *Arabidopsis thaliana*). In summary, these results show a similar contribution of the prokaryotic and fungal diversity to the structure of seed microbiota, with the exception of a few plant species that appeared dominated by prokaryotes.

**Figure 3:**
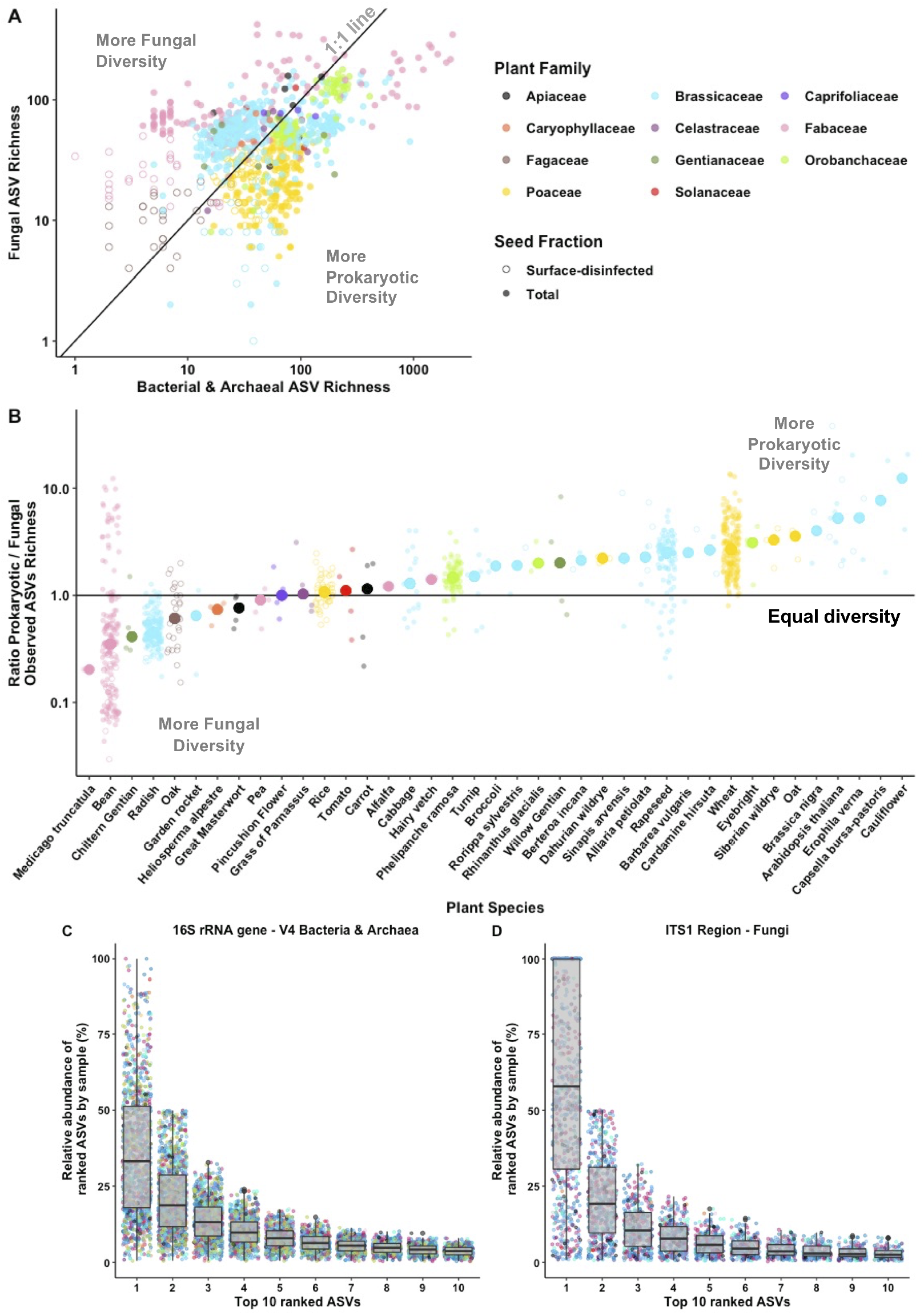
Proportion of prokaryotic (bacteria and archaea) and fungal richness in seed microbiota (n=884). A) Relationship between ASV richness values of the prokaryotic community (16S rRNA gene - V4 dataset) and of the fungal community (ITS1 region dataset). The 1 to 1 line is represented to facilitate the visualization of plant families presenting an enrichment of one of the taxonomic groups over another. Note that on panel A, both axes are on a log scale. B) Ratios of prokaryotic and fungal ASV richness for each plant species. The vertical line represents a ratio of 1 when communities present an equal diversity. Note that the y axis is presented on a log scale. Relative abundance of the top 10 ranked ASVs by sample for the prokaryotic community (C, n=1182 samples) and fungal community (D, n=850 samples). Each point represents a sample and is colored by the plant species (see legend in **Fig. 2**).

Additionally, we explored the distribution of ASVs’ relative abundance by sample using ranked abundance curves and compared the patterns between prokarytic and fungal communities (**Fig. 3C, 3D**). Both communities appeared to be dominated by few ASVs that collectively represented more than 50 or 75% of the reads in seed microbiota. In particular, we observed that seed fungal communities were generally highly dominated by one ASV (median rank 1 ASV: 57.9% relative abundance), while the rank 1 ASV of the prokaryotic community generally had a lower abundance (median: 33.1 %). It should be noted that the identity of the ranked ASVs is generally different between samples. These results indicates that seed communities are often highly uneven with few dominant prokaryotic and fungal ASVs.

#### b) Identification of six microbial phyla dominating seed microbiota

Abundance-occupancy curves were plotted for each phylum to identify the ASVs’ distribution across all samples of the meta-analysis (**Fig. 4**, **Fig. S9, S10**). The abundance-occupancy curves by phylum presented expected distributions with positive relationships between ASV abundance and prevalence, indicating that the most prevalent ASVs were also dominant in terms of relative abundance across the entire dataset. For Prokaryotes, ASVs were distributed across 43 phyla (2 archaeal and 41 bacterial phyla, **Fig. S6**) but 12 phyla were dominant in terms of relative abundance (**Fig. 4A**) and collectively represented 99.79% of the reads in the dataset and 97% of the ASV richness (7948 ASVs). In particular, the four bacterial phyla that dominate the seed microbiota both in terms of diversity and abundance were Proteobacteria, Actinobacteria, Firmicutes and Bacteroidetes (**Fig. 4A** and **Fig. S10**). A low detection of Archaea on seeds was observed with 51 ASVs that represented only 0.1% of the reads (16S rRNA gene-V4 dataset), but it should be noted that the primers targeting the 16S rRNA gene V4 region are not adapted to accurately capture archaeal diversity (Taffner et al. 2020).

**Figure 4:**
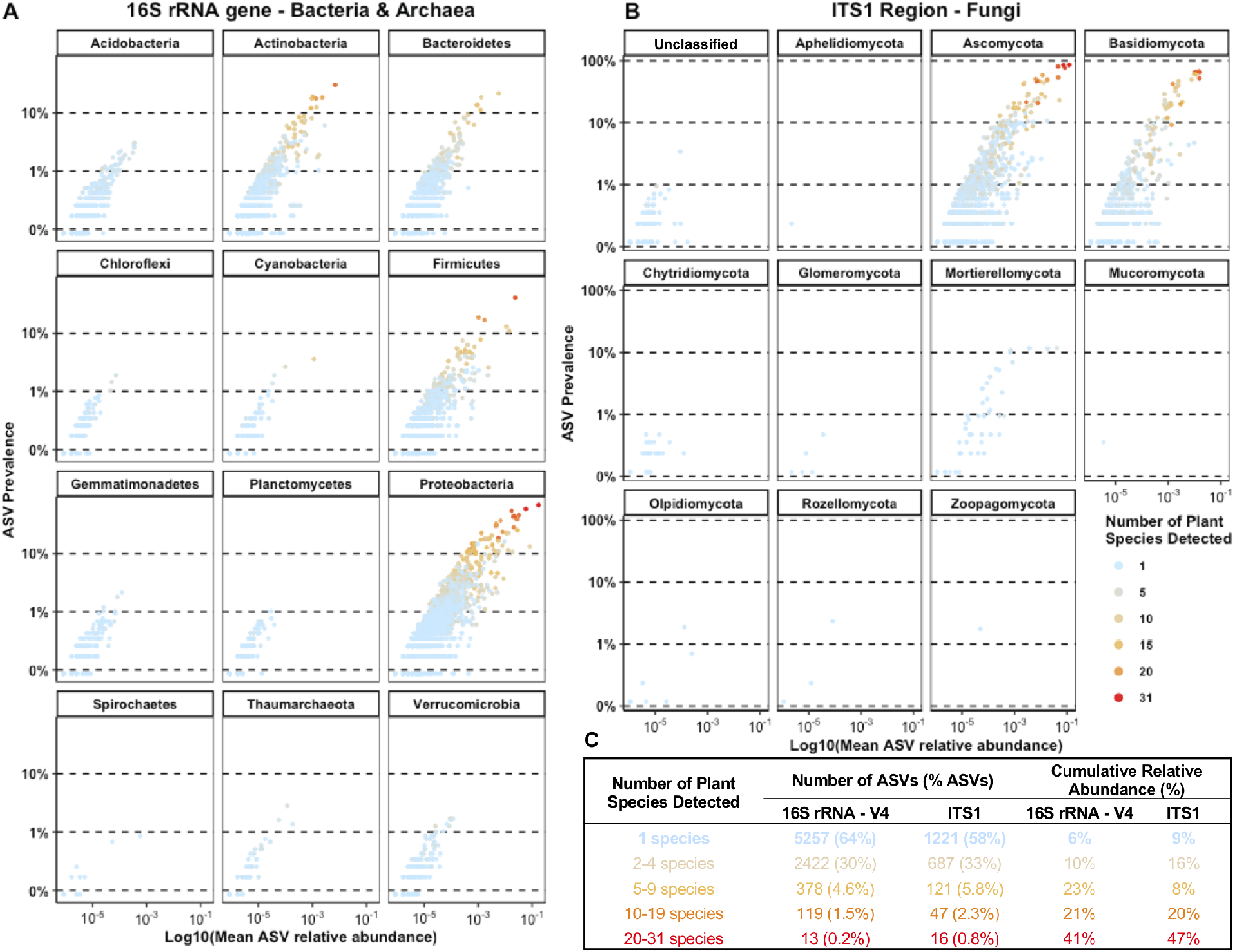
Abundance-occupancy curves for A) the most abundant prokaryotic phyla (Bacteria and Archaea, 16S rRNA gene - V4 dataset) representing more than 0.5% of the reads in the dataset and B) all fungal phyla (ITS1 region dataset). Each point represents an ASV and the points are colored based on the number of plant species in which they were detected. C) Summary table presenting the number and percentage of ASVs detected in 1 to 20-27 plant species. Note that on panels A and B, both axes are presented on a log scale.

For Fungi, ASVs were distributed across 10 phyla with two highly dominant phyla: Ascomycota (79% of reads, 61% of ASVs) and Basidiomycota (20% of reads, 31% of ASVs) that collectively represented 99.77% of the reads in the dataset and 92% of the ASV richness (7458 ASVs, **Fig. 4B**).

For each ASV, we calculated the number of plant species in which they were detected to identify highly prevalent taxa that could be considered as seed core taxa (**Fig. 4**). The vast majority of ASVs were detected in only one plant species (64% for prokaryotic ASVs and 58% for fungal ASVs), were generally detected in a few samples and not abundant (i.e. rare taxa, **Fig. 4A,B,C**). On the contrary, 13 prokaryotic ASVs and 16 fungal ASVs were detected in more than 20 different plant species and they were all extremely abundant in seed microbiota, presenting collectively a relative abundance of 41% and 47% (**Fig. 4A,B,C**). Based on the arbitrary criteria of a detection in at least 20 plant species, these taxa could be considered as core taxa as they are both prevalent and abundant in seed microbiota from diverse plants and countries.

In summary, six microbial phyla (Actinobacteria, Bacteroidetes, Firmicutes, Proteobacteria, Ascomycota and Basidiomycota) dominate seed microbiota and these communities appear to be structured in two main fractions: a core microbiota composed of few ASVs that are abundant and present in most samples (i.e. dominant taxa), and a flexible microbiota composed of a high diversity of ASVs but with an extremely variable composition between samples and a low abundance (i.e. rare taxa).

#### c) Identification of a seed core microbiota across all plant species

Next, we characterized the taxonomy, prevalence and relative abundance of each seed core ASV across all samples (**Fig. 5**). In the 16S-V4 dataset, the 13 core bacterial ASVs were predominantly Proteobacteria (n=11, **Fig. 5A**). One core bacterial ASV affiliated to *Pantoea* was highly dominant (16.9% of all reads) and was detected in 68.5% of samples collected from 27 plant species (**Fig. 5A**). The other core bacterial ASVs were affiliated to *Pseudomonas* (n=5), *Rhizobium* (n=2), *Sphingomonas, Methylobacterium* and *Paenibacillus*. The 16S rRNA gene markers permitted to obtain a taxonomic affiliation only at the genus-level but the *gyrB* marker enabled the refinement of the taxonomy of core taxa at the species level. For the *gyrB* dataset, we identified 18 bacterial ASVs present in more than 12 plant species (lower plant diversity available than in the 16S rRNA gene dataset, **Fig. 5B**, **Fig. 1B**).

**Figure 5:**
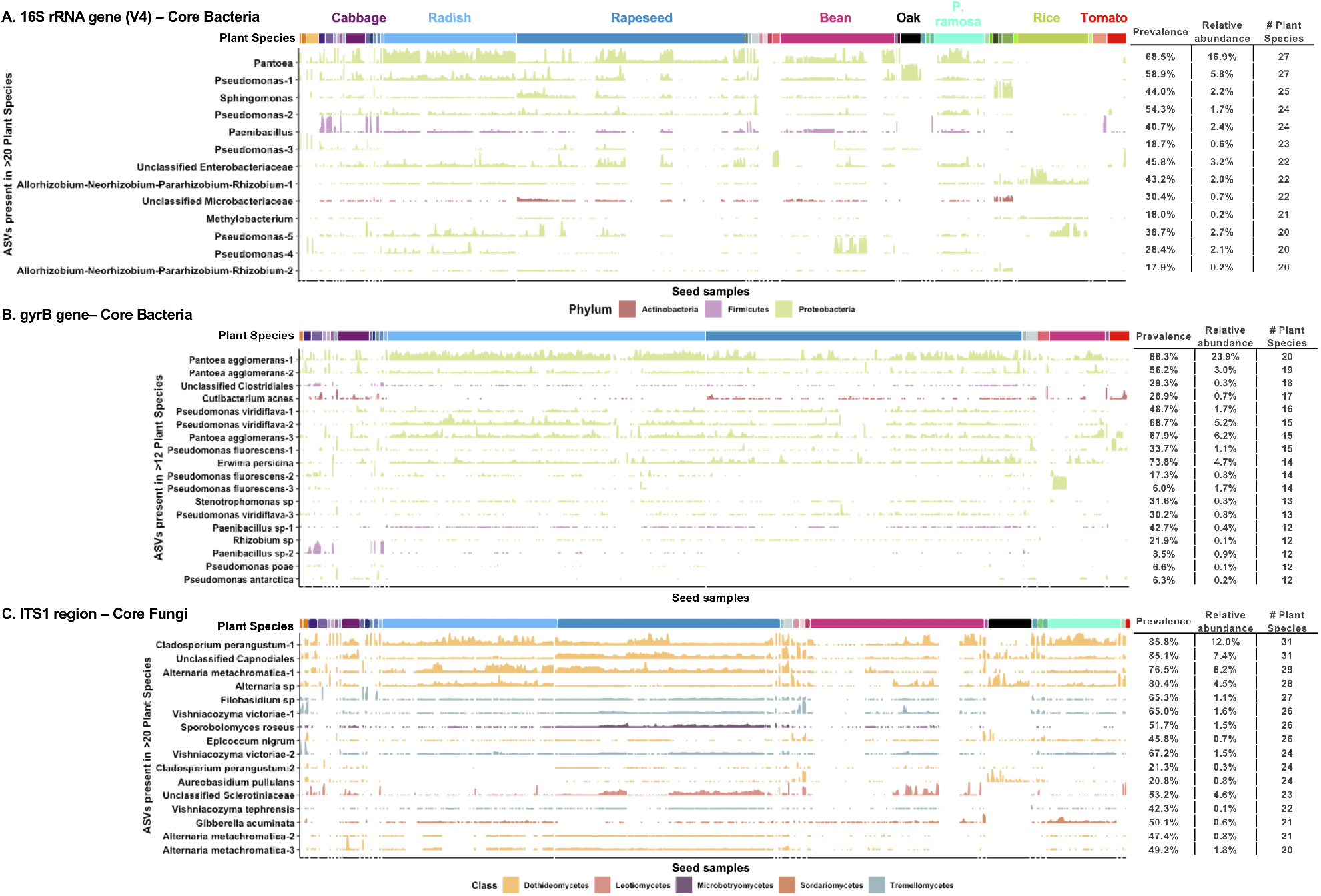
Identification of bacterial and fungal core taxa of seed microbiota across all plant species. Distribution and relative abundance of ASVs present in more than 20 plant species (>12 species for the *gyrB* dataset) across all seed samples of the meta-analysis. A) For the v4 region of 16S dataset, 13 bacterial ASVs were identified as core taxa, B) for the *gyrB* dataset, 18 bacterial ASVs were identified, and C) for the ITS1 region dataset, 16 fungal ASVs were identified. For each core ASV identified, the prevalence (% of samples), the relative abundance in the dataset (% of reads) and number of plant species (# plant species) in which the ASV is detected are presented in a table. The ridges represent the relative abundance of each ASV in the different samples of the dataset. The ASVs are ordered based on their prevalence (most prevalent ASV on top).

Therefore, the dominant *Pantoea* ASV was affiliated to the *P. agglomerans* species and was composed of three distinct ASVs, representing different populations. Interestingly, one of the three *P. agglomerans* ASVs was extremely dominant with a prevalence of 88% and a relative abundance of 24% (**Fig. 5B**). The *gyrB* dataset also indicated that *Pseudomonas* core ASVs were affiliated to four distinct species: *Pseudomonas viridiflava* (3 ASVs) and three species belonging to the *P. fluorescens* subgroup^49^: *P. fluorescens* (3 ASVs), *P*. *poae* and *P. antarctica*. Additional core taxa were identified in the *gyrB* dataset that were not detected in the 16S dataset, such as ASVs affiliated to *Cutibacterium acnes, Erwinia persicina* and *Stenotrophomonas* sp. (**Fig. 5A, 5B**). In both 16S and *gyrB* datasets, we observed that core ASVs affiliated to *Rhizobium* were present in a high number of samples but at a low abundance (except in rice).

In the ITS1 dataset, we identified 16 core fungal ASVs affiliated to the *Dothideomycetes* (9 ASVs), *Tremellomycetes* (4 ASVs), *Microbotryomycetes* (1 ASV), *Sordariomycetes* (1 ASV) and *Leotiomycetes* (1 ASV) classes (**Fig. 5C**). The core fungal community was dominated by four extremely prevalent ASVs (>77% of samples, presence in >28 plant species) with a *Cladosporium perangustum* ASV being the most abundant (12% of reads), followed by an *Alternaria metachromatica* ASV (8.2%), an unclassified *Capnodiales* ASV (7.4%) and an *Alternaria sp* (4.5%). The other core fungal ASVs were affiliated to diverse genera: *Filobasidium, Vishniacozyma, Sporobolomycetes, Epicoccum, Aureobasidium, Gibberella* (**Fig. 5C**). The full list of core and flexible taxa across all plant species is available in Supplementary file 2.

Our results suggest that the seed core microbiota is less variable on the fungal side than on the bacterial side. Based on the threshold of a detection in 20 plant species, the core fungal ASVs presented a highest prevalence (range 49-85% for fungi and 18-69% for 16S Bacteria) and were detected in a higher number of plant species (31 out of 32 for fungi and 27 out of 43 for 16S Bacteria). The higher variability in the distribution of core bacterial ASVs can be visualized in the ridge plots representing the relative abundance of each ASV in all the samples of the dataset (**Fig. 5A vs 5C**).

### Part 2: Specific patterns associated to each plant species

#### a) Plant-specific patterns in community composition

To explore the influence of plant species on seed microbiota composition, we performed ordinations based on Bray-Curtis dissimilarities between samples. On the 16S-V4 dataset, plant species were not a significant driver of seed prokaryotic community composition (**Fig. 6A**). Indeed, high variation in community composition was observed for seeds harvested from the same plant species (e.g. bean or rapeseed). In contrast, a clustering of seed microbiota per plant species was observed with ITS1. For instance, a separation between bean and rapeseed samples can be observed along the PCoA axis 1 and these two species are separated from radish samples along the PCoA axis 2. Still in many cases, an important overlap was observed between samples from different plant species, suggesting that they share similar community composition. For the 16S-V4 and *gyrB* datasets, we also performed metagenome predictions based on KEGG orthologs (KOs) using PICRUSt2 to determine if we could observe a structuring of seed microbiota based on their putative functional profiles. These metagenome predictions confirmed the limited effect of plant species on seed microbial community composition data with no clear clustering of samples by plant species and a large variability in functional profiles between samples (**Fig. S11**).

**Figure 6:**
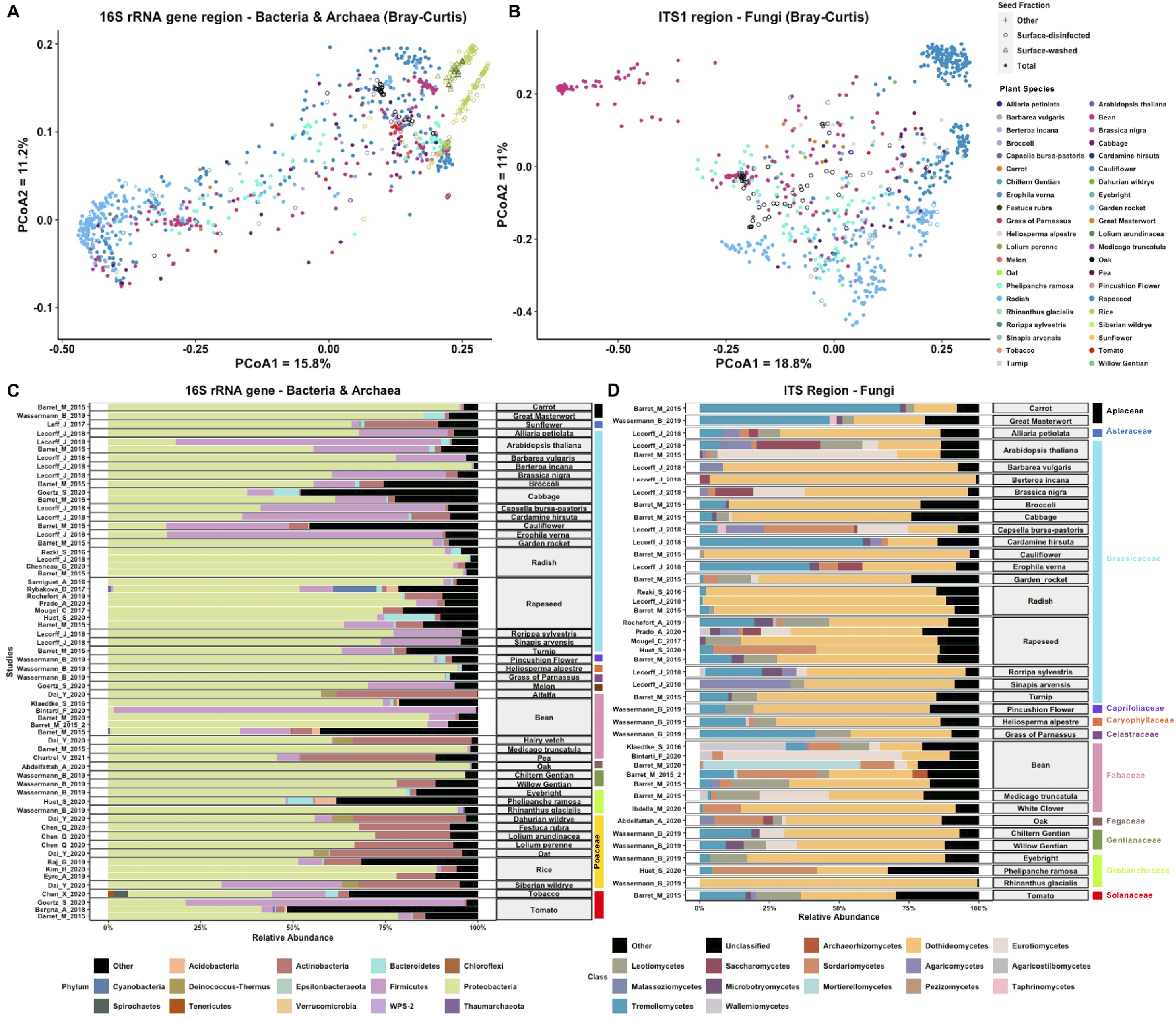
Plant-specific patterns in seed microbiota composition. Ordinations to explore the influence of plant species on seed community structure on A) the 16S rRNA gene-V4 dataset and B) the ITS1 region dataset. Average relative abundance of C) major bacterial phyla (16S-V4 dataset) and D) fungal classes (ITS1 dataset) by study for the different plant species. Species are ordered by plant family from Apiaceae at the top to Solanaceae at the bottom.

We next explored if specific patterns at the phylum or class levels existed by plant species (**Fig. 6C,D**). At the bacterial phylum level, we observed that Proteobacteria were dominant in the majority of plant species, with the exception of several Brassicaceae species that had a high abundance of Firmicutes (e.g. *Arabidopsis thaliana, Erophila verna*, **Fig. 6C**). Actinobacteria also represented a high proportion of the seed microbiota of Poaceae plants and of some Fabaceae species. Most fungal communities were dominated by taxa from the Dothideomycetes class, followed by Tremellomycetes that were present in almost all plant species with sometimes a high relative abundance (e.g. carrot, *Cardamine hirsuta*, grass of Parnassus, **Fig. 6D**). The differences observed in fungal microbiota structure between bean and rapeseed were driven by a lower abundance of Dothideomycetes in bean, which harbored a higher proportion of diverse fungal classes. The separation of these plant species with radish seed profiles was explained by a very high relative abundance of Dothideomycetes and a lower diversity of fungal classes in radish.

#### b) Identification of a seed core and flexible microbiota specific to each plant species

Next, we identified the seed core and flexible microbiota specific to some plant species for which a sufficient number of independent studies (n ≥ 3) and of samples (n ≥ 50) were available for a robust analysis. Only four plant species responded to these criteria for the 16S rRNA gene-V4 dataset (bean, radish, rapeseed and rice) and three plant species for the *gyrB* and ITS1 datasets (bean, radish, rapeseed). To identify the core taxa specific to these plant species, we used the arbitrary criteria that an ASV should be detected in a minimum of 2 independent studies and be present in at least half of the samples (minimum prevalence of 50%). Consistent with the core taxa analysis across all samples (Part 1, **Fig. 4**), we found that the core taxa identified were both extremely prevalent and abundant on seeds as this can be seen on abundance-occupancy curves (**Fig. 7A, B, C**). The core bacterial microbiota identified with the 16S rRNA gene-V4 dataset was composed of a small number of ASVs ranging from 6 ASVs for bean to 15 for radish (**Fig. 7D**). Still, these few ASVs collectively represented a large fraction of the entire bacterial community with a cumulative relative abundance of 28% (bean) to 85% (radish, **Fig. 7D**). The four plant species shared a similar core microbiota affiliated to the *Pantoea* and *Pseudomonas* genera but some core ASVs were plant-specific, like *Bacillus* for bean, *Sphingomonas* for rapeseed and rice, or *Rhizobium* and *Xanthomonas* for rice and radish (**Fig. 7G**). The core bacterial microbiota identified with the *gyrB* dataset presented similar patterns with the 16S dataset (**Fig. 7B, E, H**). The higher taxonomic resolution of the *gyrB* marker enabled to identify at the species level the core ASVs (**Fig. 7H**). The core bacterial microbiota was dominated by two *Pantoea agglomerans* ASVs and *Erwinia persicina* (not identified in 16S dataset) for the three plant species. Rapeseed and radish seeds possessed also two core ASVs affiliated to *Pseudomonas viridiflava*. Two *Pseudomonas fluorescens* core ASVs and a *Serratia marcescens* ASV were specific of rapeseed plants, while radish plants exhibited two specific *Pantoea agglomerans* ASVs and a *Pseudomonas syringae* ASV. The core fungal microbiota identified with the ITS1 dataset was generally more diverse than its bacterial counterpart, with 9 ASVs (bean) to 30 ASVs (rapeseed, **Fig. 7C, F**). These core ASVs represented an important fraction of the fungal community with a cumulative relative abundance of 35% (bean) to 87% (rapeseed, **Fig. 7F**). The three plant species shared a similar core microbiota affiliated to *Cladosporium* but many core ASVs were plant-specific, like *Mortierella* or *Fusarium* for bean, *Alternaria* or *Filobasidium* for rapeseed and radish (**Fig. 7I**).

**Figure 7:**
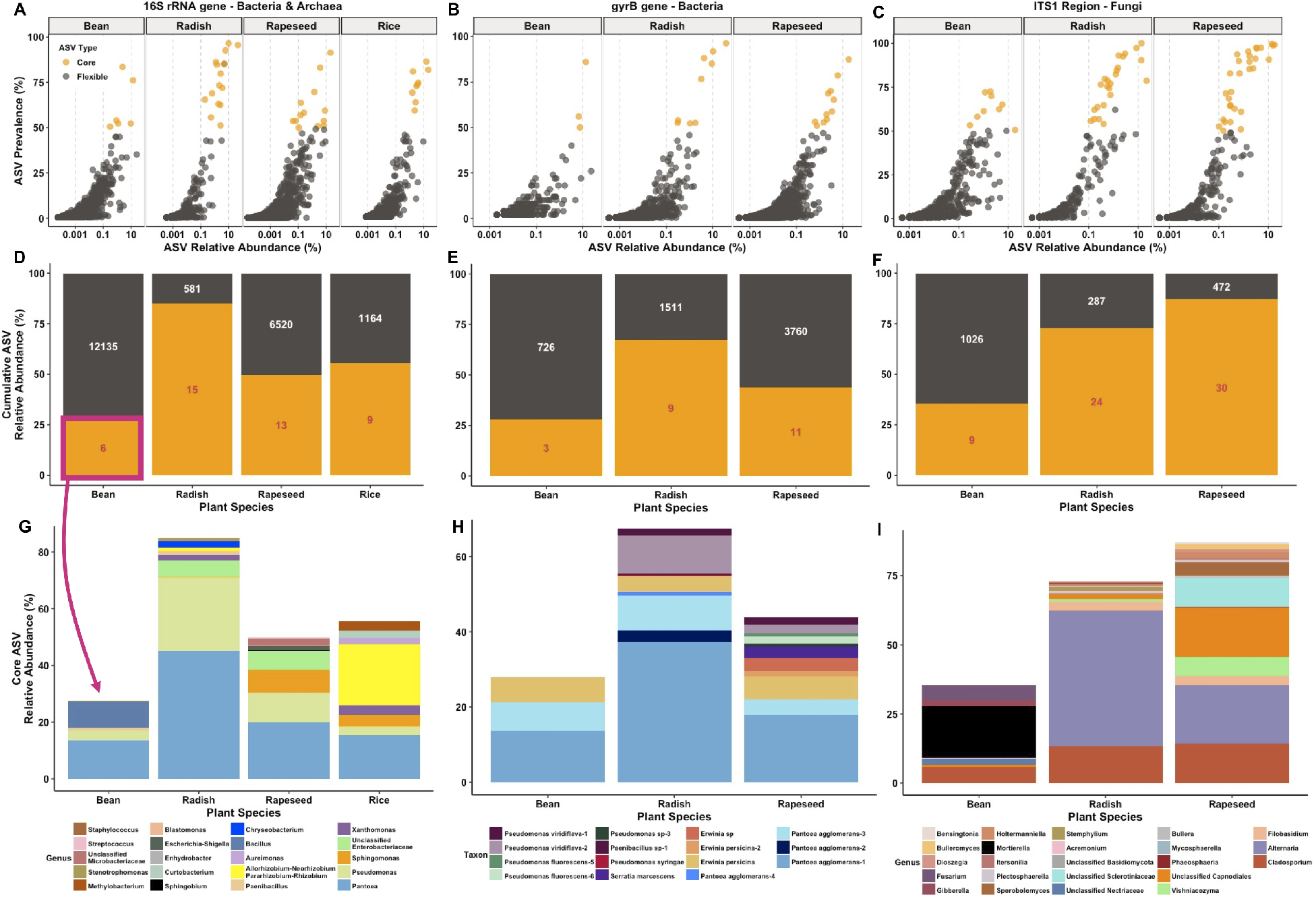
Core and flexible seed microbiota by plant species. The top panels (A, B. C) represent the abundance-occupancy curves of all ASVs by plant species for the three different marker genes (left: 16S rRNA-V4 gene, middle: *gyrB*, right: ITS1 region). This analysis was performed on 4 species for the 16S-V4 dataset and 3 species for the other two datasets based on a criteria of 3 independent studies available. Core ASVs (i.e. detection in a minimum of 2 studies and a minimum 50% prevalence) are highlighted in yellow. The middle panels (D, E, F) display the cumulative relative abundance of core and flexible ASVs by plant species. The number inside each bar indicates the number of ASVs for each group (in red for core ASVs, in white for flexible ASVs). The bottom panels (G, H, I) represent the taxonomic composition of the core microbiota of each plant species, at the genus level for the 16S-V4 and ITS1 datasets and at the species level for the gyrB dataset.

This analysis also enabled to characterize the diversity and relative abundance of the flexible seed microbiota that varied widely between plant species. Bean seeds presented the most flexible microbiota representing more than 65% of the relative abundance and a high ASV diversity (12135 ASVs for 16S-V4 and 1026 ASVs for ITS1). Radish seeds had the least flexible microbiota, likely because most of the studies available originated from our research group and a single variety (Flamboyant5). The full list of core and flexible taxa by plant species is available in Supplementary file 2.

These results confirm that seed microbiota share many similarities across plant species, especially on the bacterial side with no clear clustering by plant species and several core taxa shared. For the fungal community, we observed more dissimilarities between plant species, especially with some compositional differences between bean, radish and rapeseed seeds.

## DISCUSSION

Seed microbiota are diverse and extremely variable in the number of microbial taxa capable of colonizing seed samples, from one to thousands of ASVs, with a median of a hundred ASVs (prokaryotes + fungi). These taxa overwhelmingly originate from four bacterial phyla (Proteobacteria, Actinobacteria, Firmicutes and Bacteroidetes) and two fungal phyla (Ascomycota and Basidiomycota), which is similar with other plant compartments^50^. However, our findings show that seeds generally present similar proportions of bacterial and fungal diversity, which contrasts with other plant compartments that are generally highly dominated by bacterial diversity ^50–52^. This meta-analysis highlights that seed microbiota present stable (ie. core) and variable (i.e flexible) microbial fractions across samples, as already hypothesized for the plant microbiome^8^ and observed in other microbial ecosystems ^53,54^. A striking result of this meta-analysis is that around 30 bacterial and fungal taxa are shared between very contrasted plant species and detected in samples from all over the world. In particular, core ASVs affiliated to the *Pantoea, Pseudomonas, Cladosporium* and *Alternaria* genera appear as extremely abundant and prevalent seed-borne microorganisms. It is worth noting that this study confirms that arbuscular mycorrhizal fungi are not frequently transmitted to seedlings through seeds, with very few Glomeromycota ASVs detected and at a very low prevalence (0.3% of seed samples)^55^. In contrast, two *Rhizobium* ASVs were identified as core ASVs across all samples with high prevalence and generally low abundances, indicating that transmission through seeds is likely a prevalent pathway for these potential nitrogen-fixing bacteria ^56^. The core taxa identified represent a short-list of microbial taxa of interest to investigate their potential role in plant fitness, such as seed maturation, germination or emergence, and to identify their environmental origin (horizontal or vertical transmission). We encourage future studies to attempt to isolate and characterize these core taxa using genomics, synthetic community reconstruction and thorough plant phenotyping to identify their adaptation to the seed habitat and their potential selection by plant hosts. Still, the majority of seed microbiota belongs to the “flexible” fraction that is reflective of the abiotic and biotic fluctuations of the seed environment. These taxa are also adapted to the seed habitat but generally harbor lower abundances and prevalence. This fraction should not be neglected as it represents an important diversity reservoir that can be transmitted to seedlings ^57,58^ and provide microbial taxa that are adapted to specific local constraints ^8,53^.

This meta-analysis provides new insights on seed microbiota diversity and composition but also highlights key knowledge gaps. This study gathered data on only 50 plant species (including 30 crops), with often only one study by plant species and few seed samples analyzed. This means that the seed microbiota of a majority of crops and natural plant communities have not been characterized. More investigations are also required to assess the influence of geographical sites and plant genotypes on seed microbiota. Additionally, this meta-analysis included mainly seed microbiota characterization from multiple seeds (>3 seeds in sample) with more than 1000 reads. More work is needed at the single seed level ^37,59,60^ to assess seed-to-seed variation and the proportion of seeds that do not have detectable microbiota (i.e. sterile seeds)^9,61^. Moreover, seeds can greatly vary in their chemical composition, size, anatomic features and physiology but we still do not know how these seed attributes influence microbiome composition. Unfortunately, the diverse data gathered in this meta-analysis cannot robustly address this question and future studies specifically designed to characterize the links between seeds traits and microbiota are needed. Another major knowledge gap in the seed microbiota literature regards the characterization of other microorganisms than bacteria and fungi, especially Oomycetes and other protists, Archaea and viruses. The most common primers used for amplicon sequencing of seed microbiota completely miss these taxonomic groups (Oomycetes, protists, viruses) or partially characterize them (Archaea), despite knowledge on their presence on seeds from cultivation-based approaches and phytopathological studies ^62–64^. To allow the Seed Microbiota Database to grow in the future, we encourage new studies to perform multi-markers analyses on a minimum of 5 replicate seed samples and to deposit their data on public databases with detailed metadata to allow reusability. These future studies will help gaining a more comprehensive view of seed microbiota and will accelerate discoveries using seeds as a vector of agricultural innovations, especially for plant microbiome engineering.

## METHODS

### Studies included in the Seed Microbiota Database

We identified seed microbiota studies from our team and from the literature by performing keyword searches in Google Scholar and by following references in reviews and seed microbiota papers (end of search November 17th, 2020). We initially identified 100 seed microbiota studies (59 for 16S rRNA gene; 14 for *gyrB* gene and 27 for ITS region) that used amplicon sequencing (i.e metabarcoding), but we decided to conserve in the meta-analysis only studies using the Illumina sequencing technology that was the most common approach (90 datasets). This choice permitted us to use a similar bioinformatic pipeline for all studies and to not introduce biases in our interpretation due to sequencing technologies. Out of these 90 studies, only 63 studies were finally included in this meta-analysis based on the following criteria: i) availability of the raw sequence data on public repository (fastq files), ii) availability of metadata associated to each sample, iii) sequencing quality, and iv) the primer sets used. For this last point, we focused on the primers or gene regions that were the most commonly studied in seed microbiota studies: for the 16S rRNA gene the V4 and V5-V6 regions and for ITS, the ITS1 and ITS2 regions. Most studies were downloaded from online repositories (e.g. ENA, NCBI-SRA) and some were obtained after personal communication with the authors. Detailed information on the 63 studies that constitute the Seed Microbiota Database can be found in Supplementary Data 1.

### Bioinformatic analysis and preparation of the datasets and subsets

Each study was independently re-processed with a standardized bioinformatic pipeline (code available at https://github.com/marie-simonin/Seed-Microbiota-Database) using Qiime 2 and DADA2^65,66^. In brief, primer sequences were removed with cutadapt 2.7 and trimmed fastq files were processed with DADA2 version 1.10, using variable truncation parameters depending on the sequencing quality of the study. If necessary, because of small differences in the primers used, a trimming parameter was added in DADA2 to have all studies targeting the exact same region of the V4, V5-V6, ITS1 or ITS2 regions. Chimeric sequences were identified and removed with the removeBimeraDenovo function of DADA2. ITS1 and ITS2 reads were processed similarly, except that only the forward reads were included in the analysis as recommended by Pauvert et al. (2020)^67^ to maximize fungal diversity detection. ASV taxonomic affiliations were performed with a scikit-learn multinomial naive Bayesian classifier implemented in Qiime 2 (qiime feature-classifier classify-sklearn^68^). ASVs derived from 16S rRNA reads were classified with the SILVA 132 database^69^ trained for the specific gene region (V4 or V5-V6). *gyrB* reads were classified with an in-house *gyrB* database (train_set_gyrB_v4.fa.gz) available upon request. ASVs derived from ITS1 and ITS2 reads were classified with the UNITE v8 fungal database^70^. Unassigned ASVs or ASVs affiliated to mitochondria, chloroplasts or eukaryotes were removed from the 16S rRNA gene studies. Non-fungal ASVs were filtered from the ITS gene studies, and unassigned and *parE* ASVs (a *gyrB* paralog) were filtered from the *gyrB* gene studies. After these filtering steps, samples with less than 1000 reads were excluded from the meta-analysis. For each study, two datasets were prepared:an unrarefied and a rarefied one. The rarefaction was performed at the lowest read number observed (min=1000 reads/sample, max=86108 reads/sample, Supplementary file 1).

The next step was to merge all the studies targeting the same marker gene and region to form a final “dataset”. This merging step between different studies was possible because all the ASVs obtained targeted the exact same region of the marker gene considered (see above). A total of 5 datasets (16S rRNA-V4, 16S rRNA-V5-V6, *gyrB*, ITS1 and ITS2) was included in the meta-analysis (**Fig. 1B**). The unrarefied studies were merged into one dataset then further filtered to obtain a “clean database” (e.g. Subset 1) by removing i) ASVs present in only one sample and with less than 20 reads in the entire dataset, ii) ASVs shorter than 200 bp, and iii) *gyrB* ASVs with a read length not compatible with codon triplets (multiple of 3). The rarefied studies were merged into one dataset (Subset 2) to look at study-specific patterns in seed microbiota diversity. To investigate community structure and composition patterns across all studies, an additional Subset was prepared by performing a global rarefaction across all samples (Subset 3, rarefaction at 1000 to 1056 reads/sample depending on the dataset, Table S1).

### Community and Taxa-level analyses

Diversity and community structure analyses were performed in R 3.6.2 using the *phyloseq* (v1.28.0), *vegan* (v2.5-7) and *microbiome* (v1.7.21) packages^71–73^. The code and files to reproduce the analyses and figures are available at https://github.com/marie-simonin/Seed-Microbiota-Database. Seed microbiota diversity across all studies was explored using ASV richness, Shannon and Pielou’s evenness indexes. Changes in seed microbial community composition were assessed using Bray-Curtis dissimilarity and principal coordinate analysis (PCoA) was used to plot the ordination.

Seed core and flexible microbiota across all plant species or specific to some plant species were identified. For the analyses across all plant species, we used the arbitrary criteria that ASVs detected in at least 20 plant species would be considered as core taxa, as they are prevalent in seed microbiota from diverse plants and countries.

For the analyses by plant species, we focused on plants for which a sufficient number of independent studies (minimum 3 studies) and samples (minimum 50 samples) was available for a robust analysis. Four plant species responded to these criteria for the 16S rRNA gene-V4 dataset (bean, radish, rapeseed and rice) and three plant species for the *gyrB* and ITS1 datasets (bean, radish, rapeseed). To identify the core ASVs specific to these plant species, we used the arbitrary criteria that an ASV should be detected in a minimum of 2 independent studies and be present in at least half of the samples (minimum prevalence of 50%). For the 16S-V4 and *gyrB* datasets (using Subset 1), we used PICRUSt2^74^ to predict metagenome functions of seed microbiota from metabarcoding data.

All figures were prepared using the *ggplot2* (v3.3.3)^75^ and *ggpubr* (v0.4.0) packages and the data management was done using the *dplyr* (v1.0.4) and *tidyverse* (v1.3.0) packages in R and in DB Browser for SQLite (v3.12.2).

## ACKNOWLEDGEMENTS

We are deeply grateful to the authors whose data contributed to this database and meta-analysis. Without their efforts to make their datasets available with good metadata, this meta-analysis would not have been possible. We thank all our collaborators that contributed unpublished datasets to this study, in particular Josiane Le Corff, Muriel Marchi and Marie-Anne Le Moigne for the Endowild project (Le Corff_J_2018 study), Simon Goertz (Goertz_S_2020 study), Christophe Mougel for the Brassica Div Patho project (Mougel_C_2017 study) and Brice Marolleaux for the Seed Microbiome project (Barret_M_2020 study).

## DATA AVAILABILITY

The Seed Microbiota Database is available on the Data INRAE portal (https://doi.org/10.15454/2ANNJM). The database includes all the metadata tables, ASV tables, taxonomy tables, fasta files and phylogenetic trees associated to the 5 datasets corresponding to the different molecular markers. Two scripts are available to query the database: 1) the “getASVInfosInDb.pl” to search the database for a specific ASV using its sequence; 2) the “getASVFoundOnSpecies.pl” to get the list of all ASVs associated with a plant species. More details are available on the README file in the repository.

The code and files to reproduce the bioinformatic analyses and figures are available at https://github.com/marie-simonin/Seed-Microbiota-Database. The list of accession numbers (e.g. NCBI-SRA, ENA) to download the raw data of the studies included in the database is available in the Supplementary file 1 - Study description.

## SUPPLEMENTARY INFORMATION

**Table S1:**
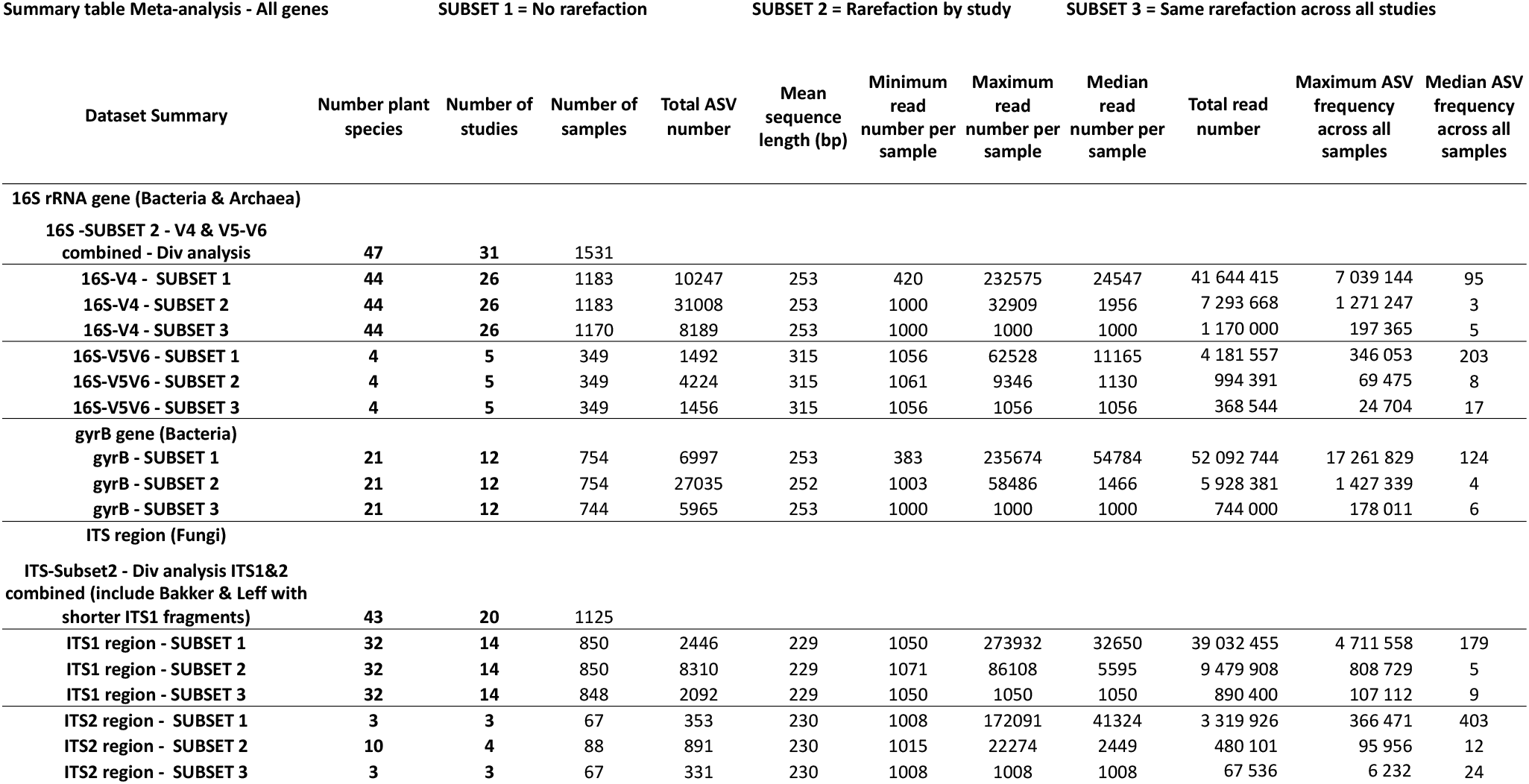
Detailed description of all the datasets and subsets used in the meta-analysis for the three marker genes.

**Figure S1:**
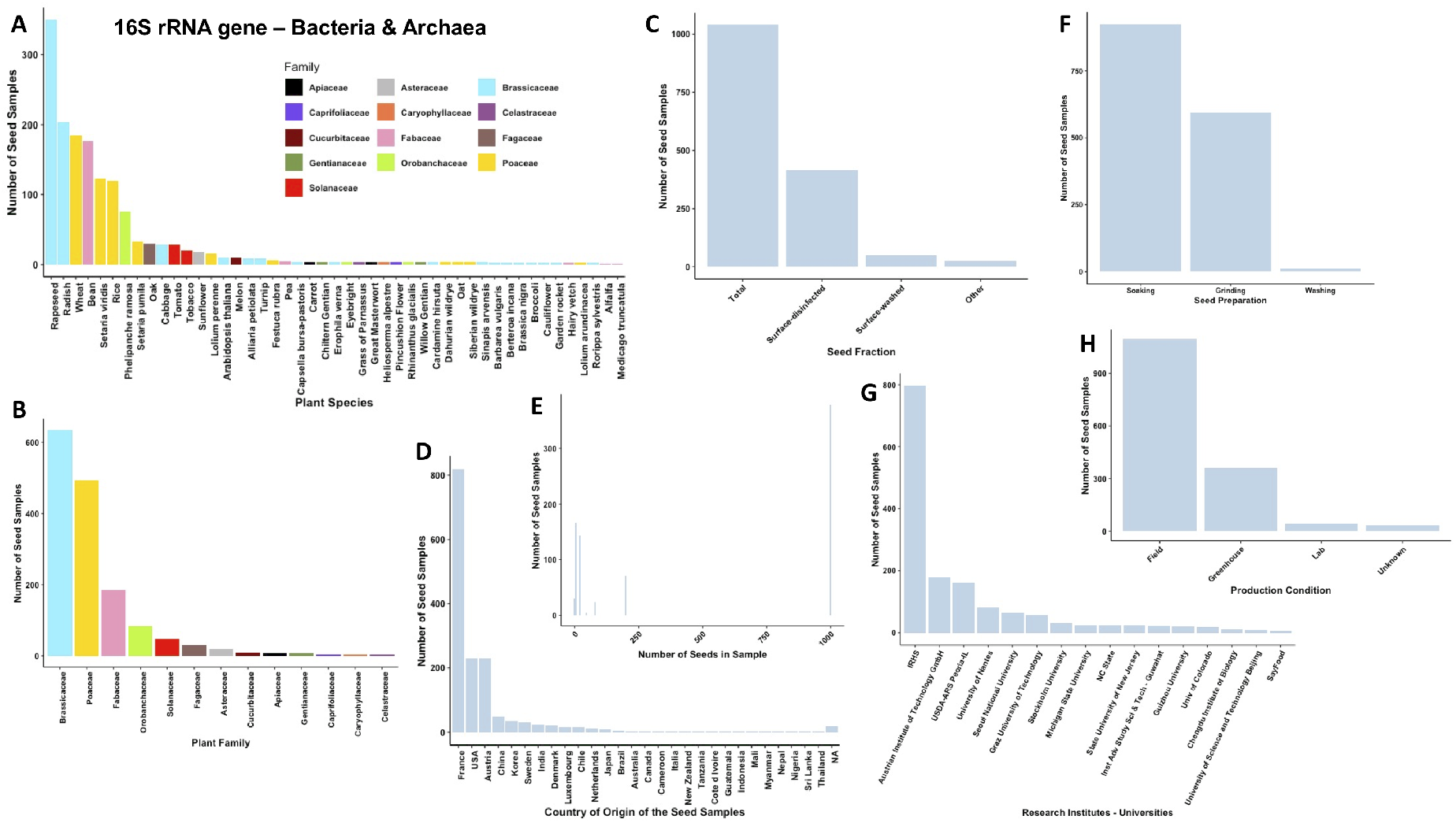
Description of the 16S rRNA gene samples (both V4 and V5-V6 regions) included in the meta-analysis. Distribution of seed samples: A) by plant species, B) by plant family, C) depending on the seed fraction considered, D) by country of origin, E) based on the number of seeds included in the sample, F) depending on the seed preparation before DNA extraction, G) by research institute or university that produced the results, H) based on the production conditions of the seed samples.

**Figure S2:**
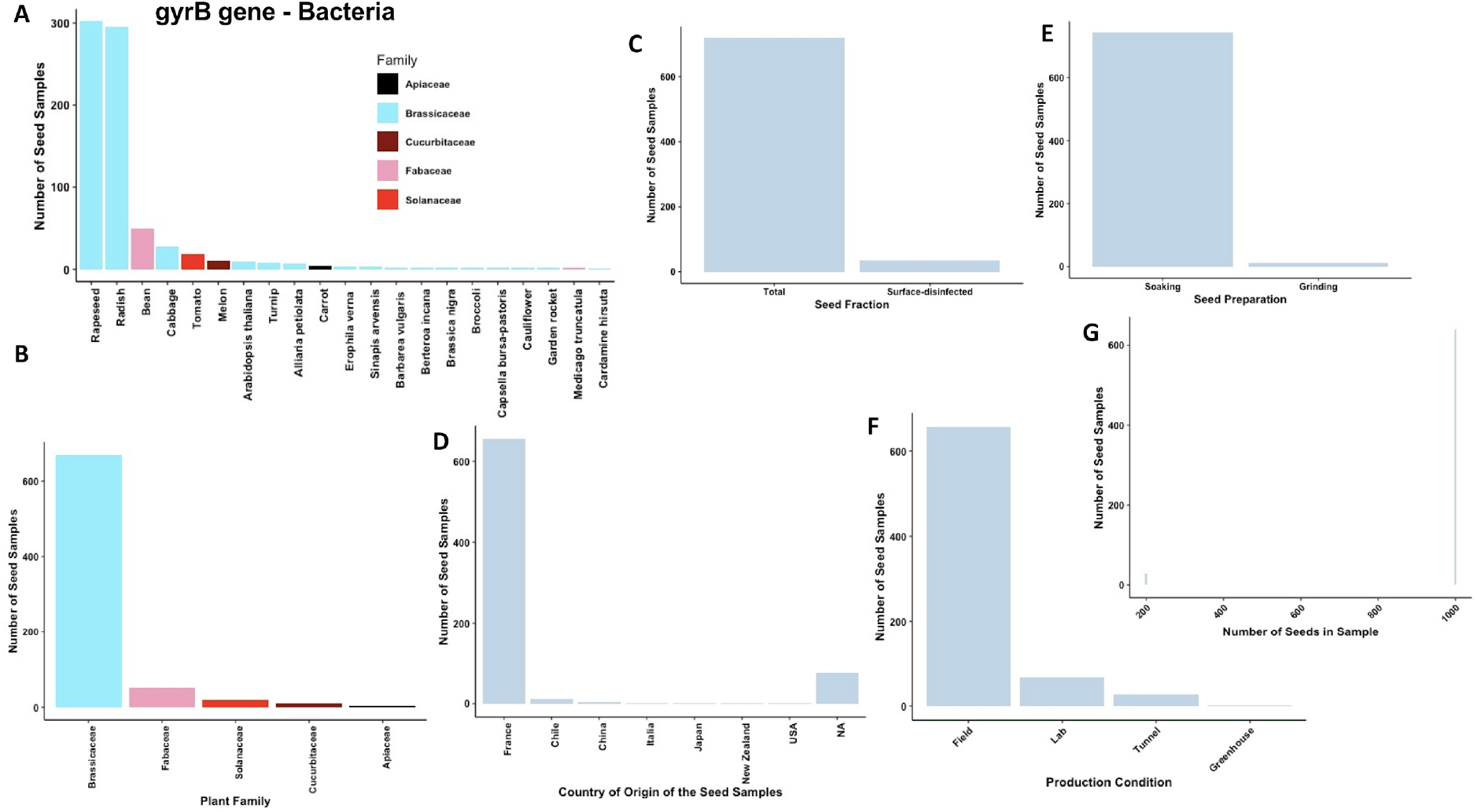
Description of the gyrB gene samples included in the meta-analysis. Distribution of seed samples: A) by plant species, B) by plant family, C) depending on the seed fraction considered, D) by country of origin, E) depending on the seed preparation before DNA extraction, F) based on the production conditions of the seed samples, G) based on the number of seeds included in the sample. No plot was displayed regarding the research institutes or universities, because all the gyrB studies were realized in our team at the IRHS.

**Figure S3:**
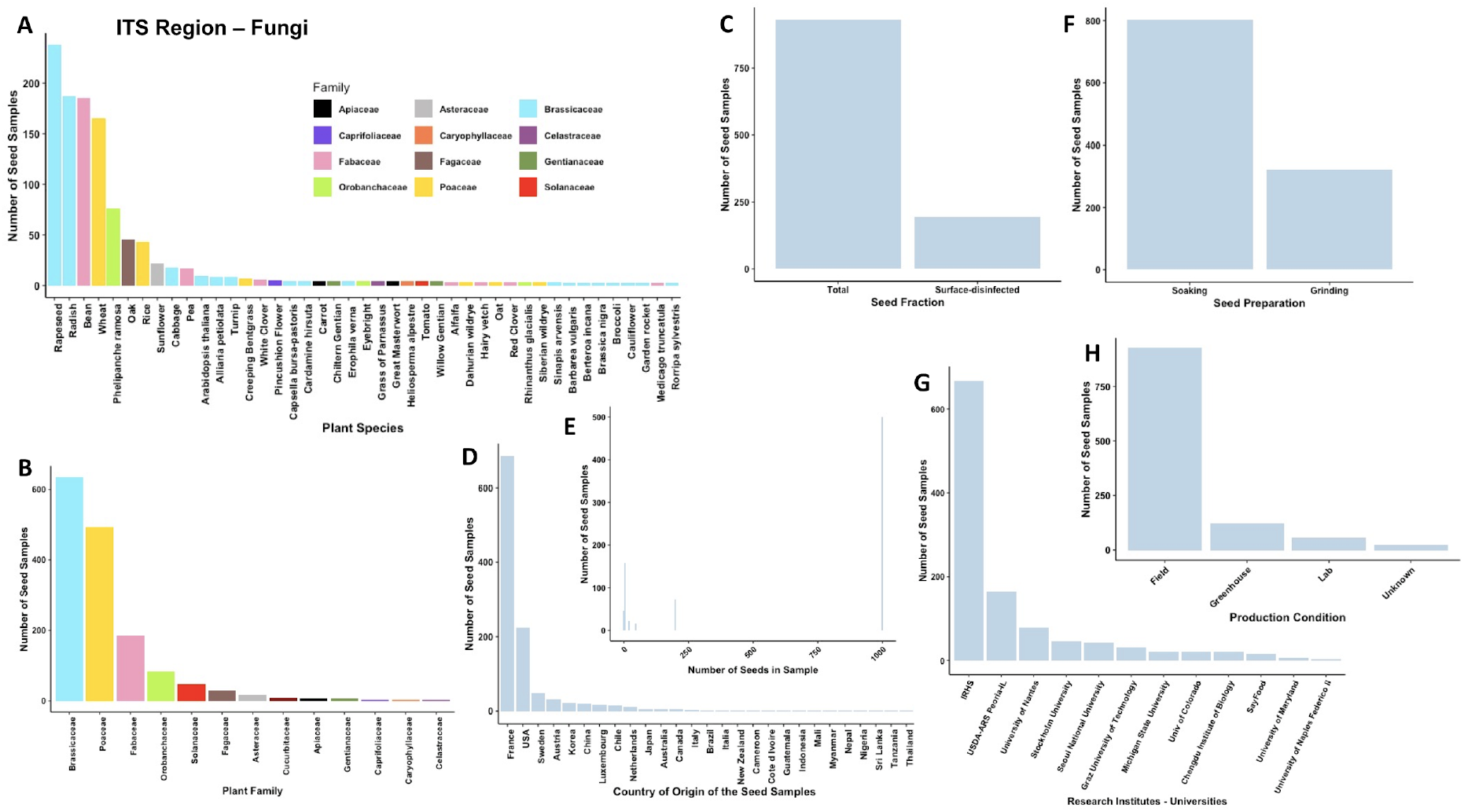
Description of the ITS samples (both ITS1 and ITS2 regions) included in the analysis. Distribution of seed samples: A) by plant species, B) by plant family, C) depending on the seed fraction considered, D) by country of origin, E) based on the number of seeds included in the sample, F) depending on the seed preparation before DNA extraction, G) by research institute or university that produced the results, H) based on the production conditions of the seed samples.

**Figure S4:**
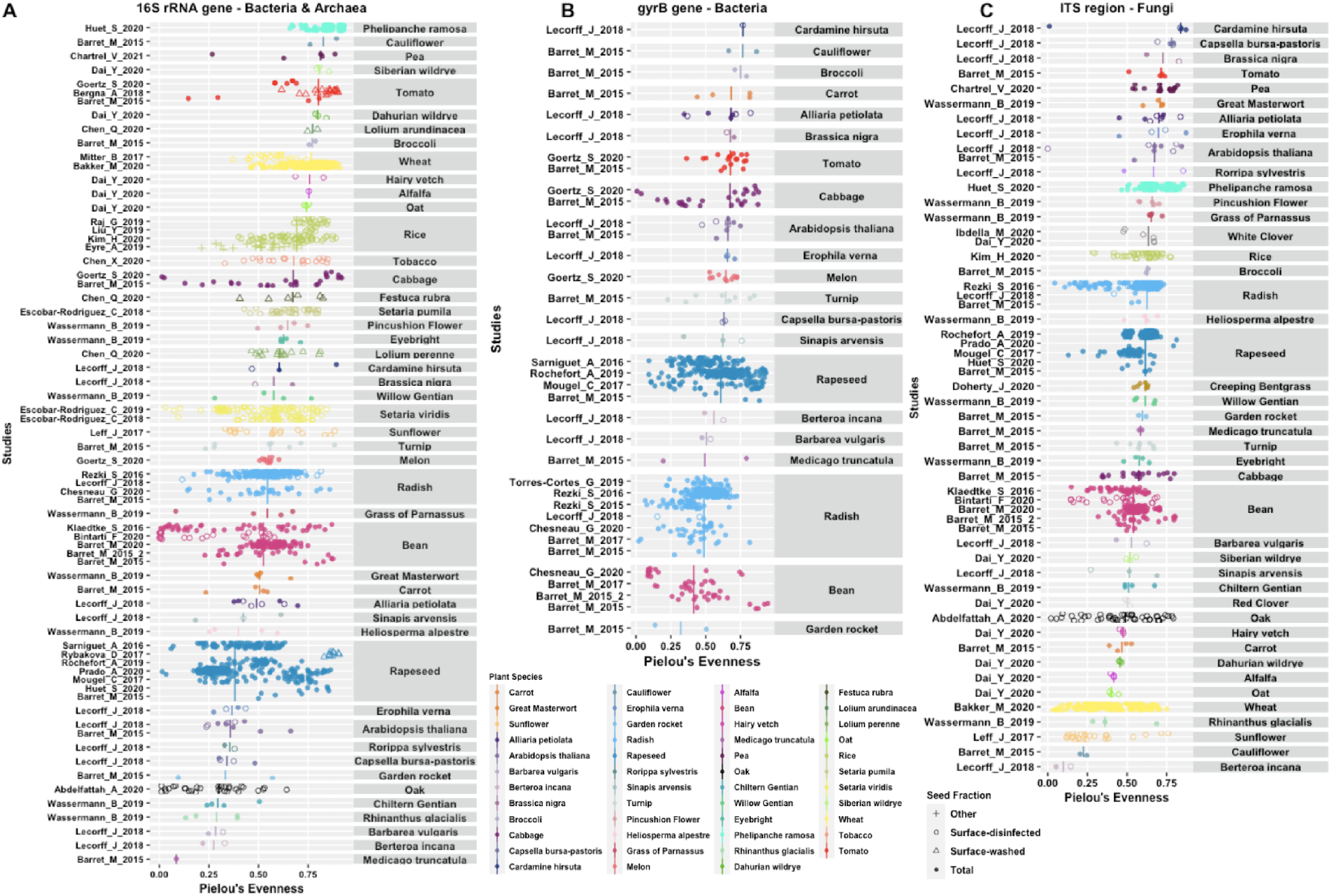
Pielou’s evenness index values of seed microbiota within and across plant species. Evenness values of all the seed samples included in the meta-analysis for the 16S rRNA marker gene (panel A, n=1531), the gyrB marker gene (panel B, n=754) and the ITS region marker (panel C, n=1125). Each point represents a seed sample that can be composed of multiple seeds (up to a thousand seeds). The shape of the points represents the sample preparation performed before DNA extraction of the seeds (Total: no pre-treatment of the seeds, Surface-washed: seeds were rinsed with sterile water, Surface-disinfected: seeds were surface sterilized with chemical products, Other: seeds were dissected or received specific treatments). For each plant species, the vertical line corresponds to the median value of the samples.

**Figure S5:**
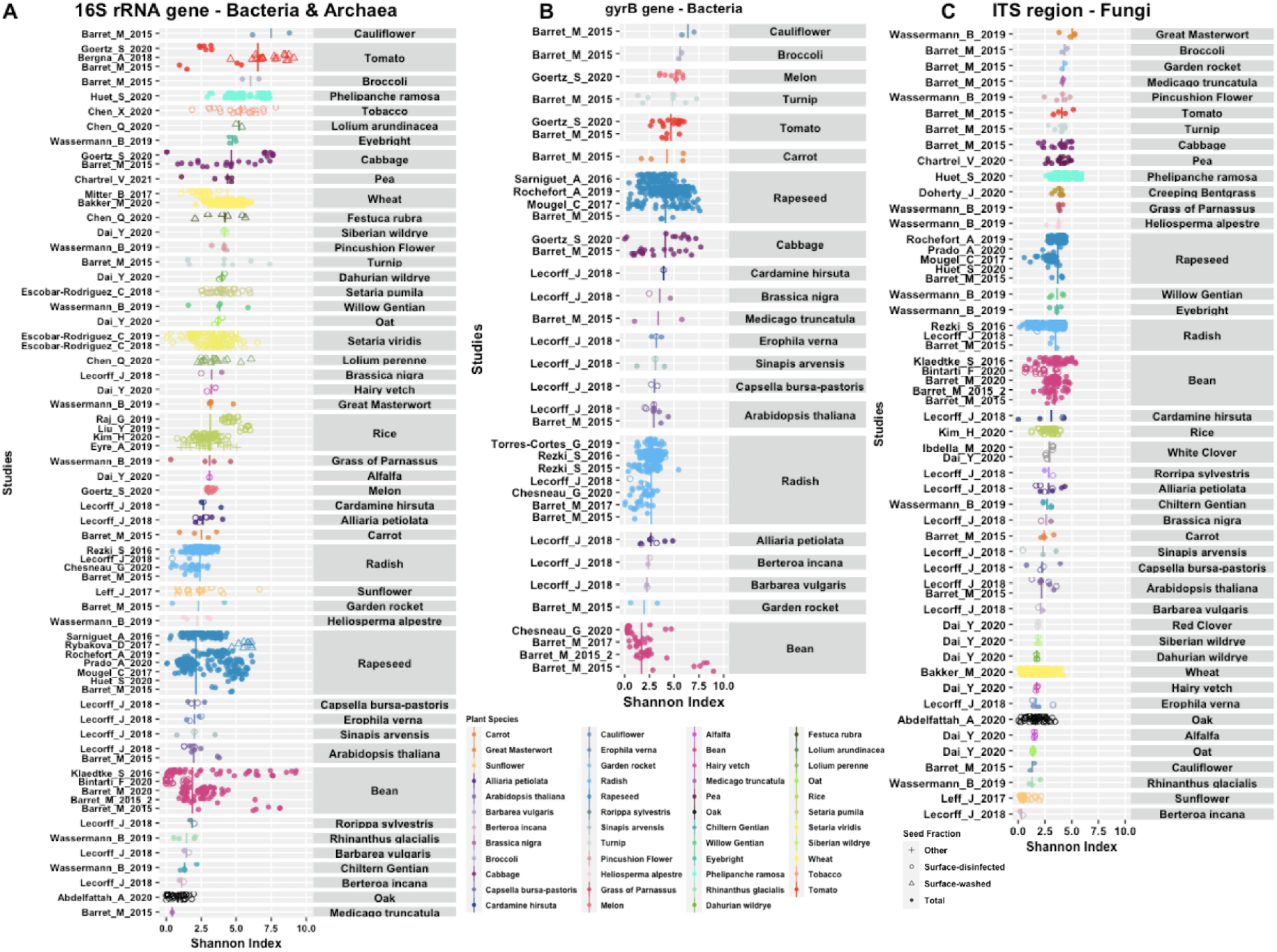
Shannon diversity index values of seed microbiota within and across plant species. Shannon index of all the seed samples included in the meta-analysis for the 16S rRNA marker gene (panel A, n=1531), the gyrB marker gene (panel B, n=754) and the ITS region marker (panel C, n=1125). Each point represents a seed sample that can be composed of multiple seeds (up to a thousand seeds). The shape of the points represents the sample preparation performed before DNA extraction of the seeds (Total: no pre-treatment of the seeds, Surface-washed: seeds were rinsed with sterile water, Surface-disinfected: seeds were surface sterilized with chemical products, Other: seeds were dissected or received specific treatments). For each plant species, the vertical line corresponds to the median value of the samples.

**Figure S6:**
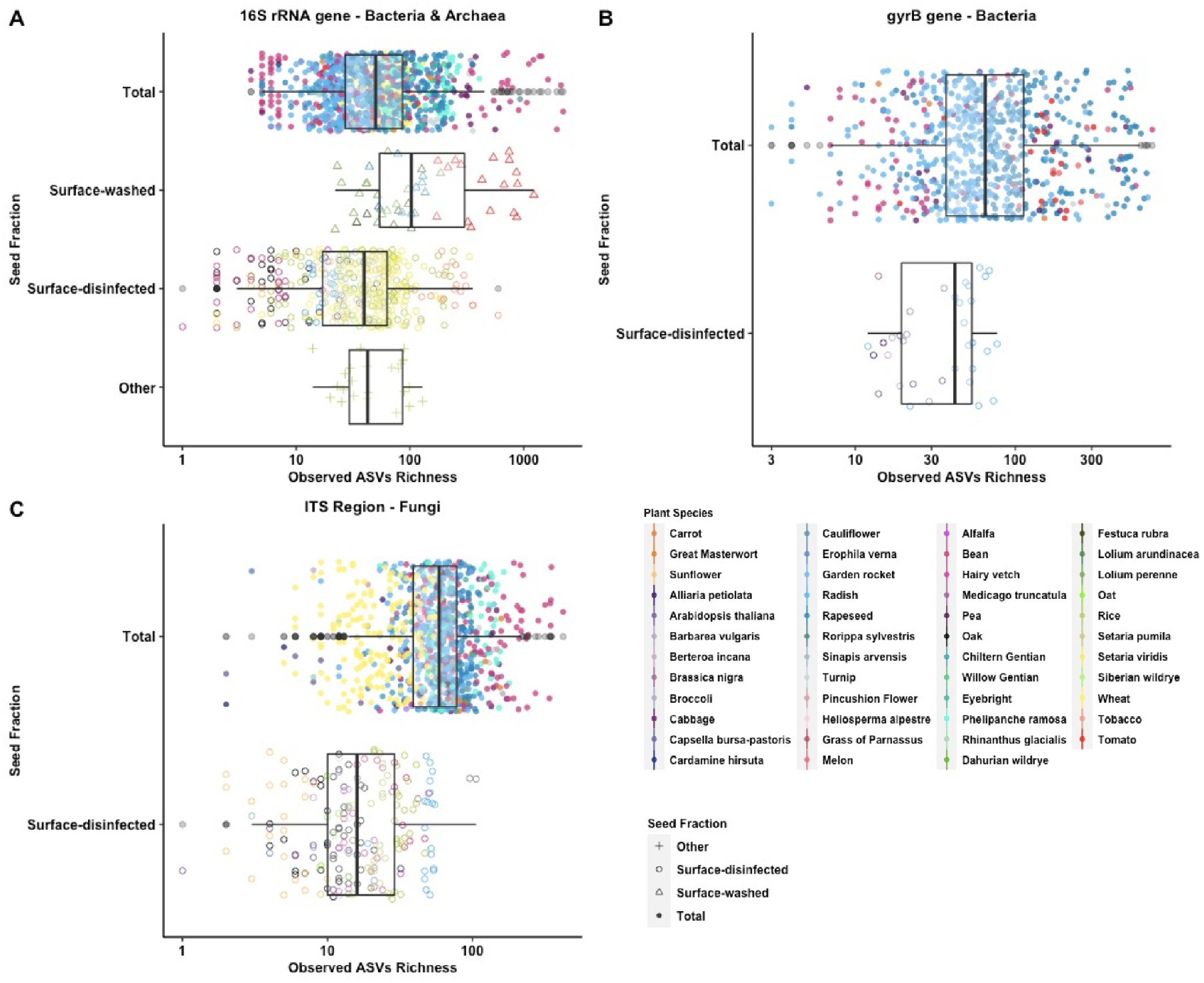
Seed microbiota diversity in function of the seed fraction considered (Total, Surface-disinfected, Surface-washed, Other). Total: no pre-treatment of the seeds, Surface-washed: seeds were rinsed with sterile water, Surface-disinfected: seeds were surface sterilized with chemical products, Other: seeds were dissected or received specific treatments.

**Figure S7:**
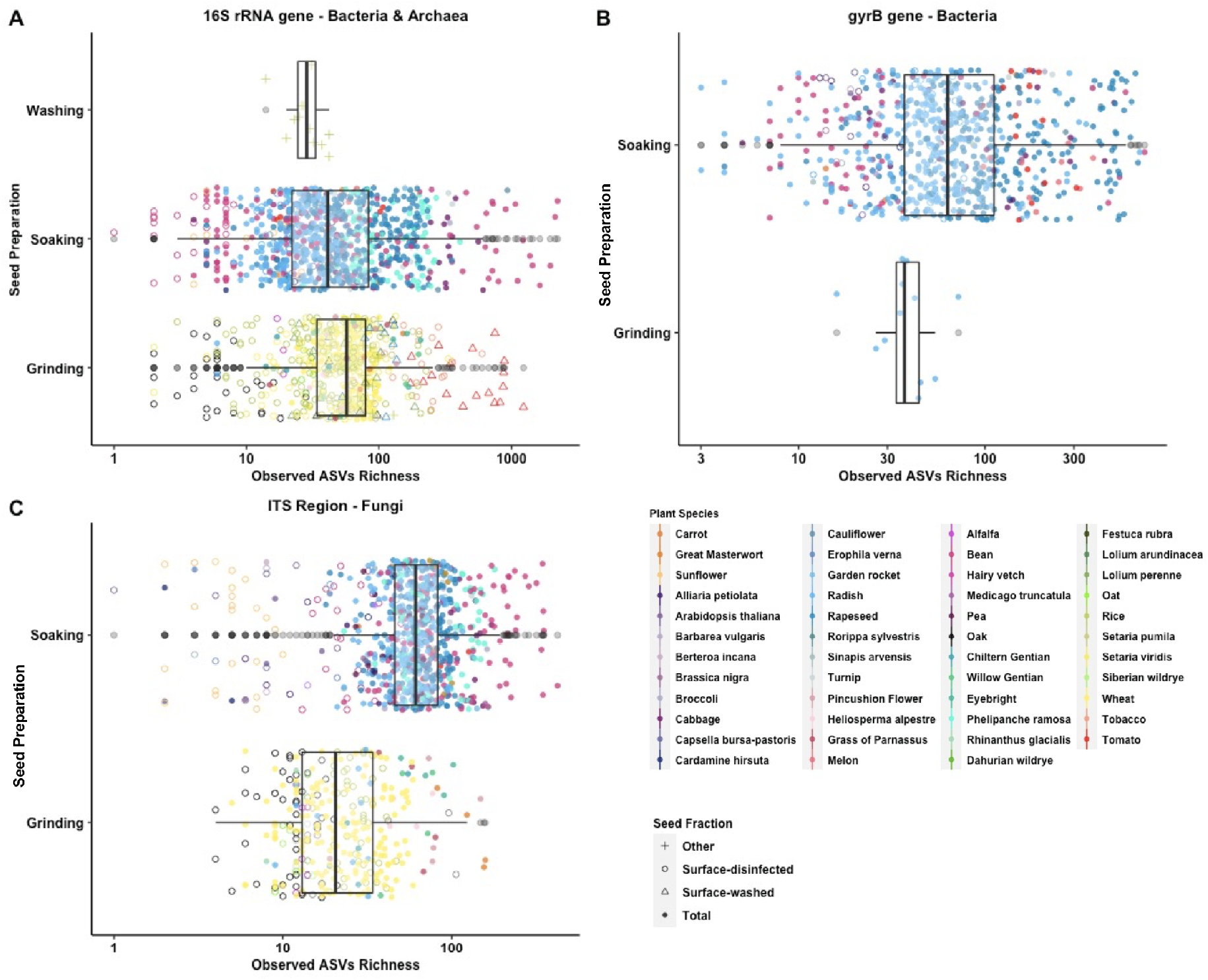
Seed microbiota diversity in function of the seed preparation before DNA extraction (Soaking, Grinding, Washing).

**Figure S8:**
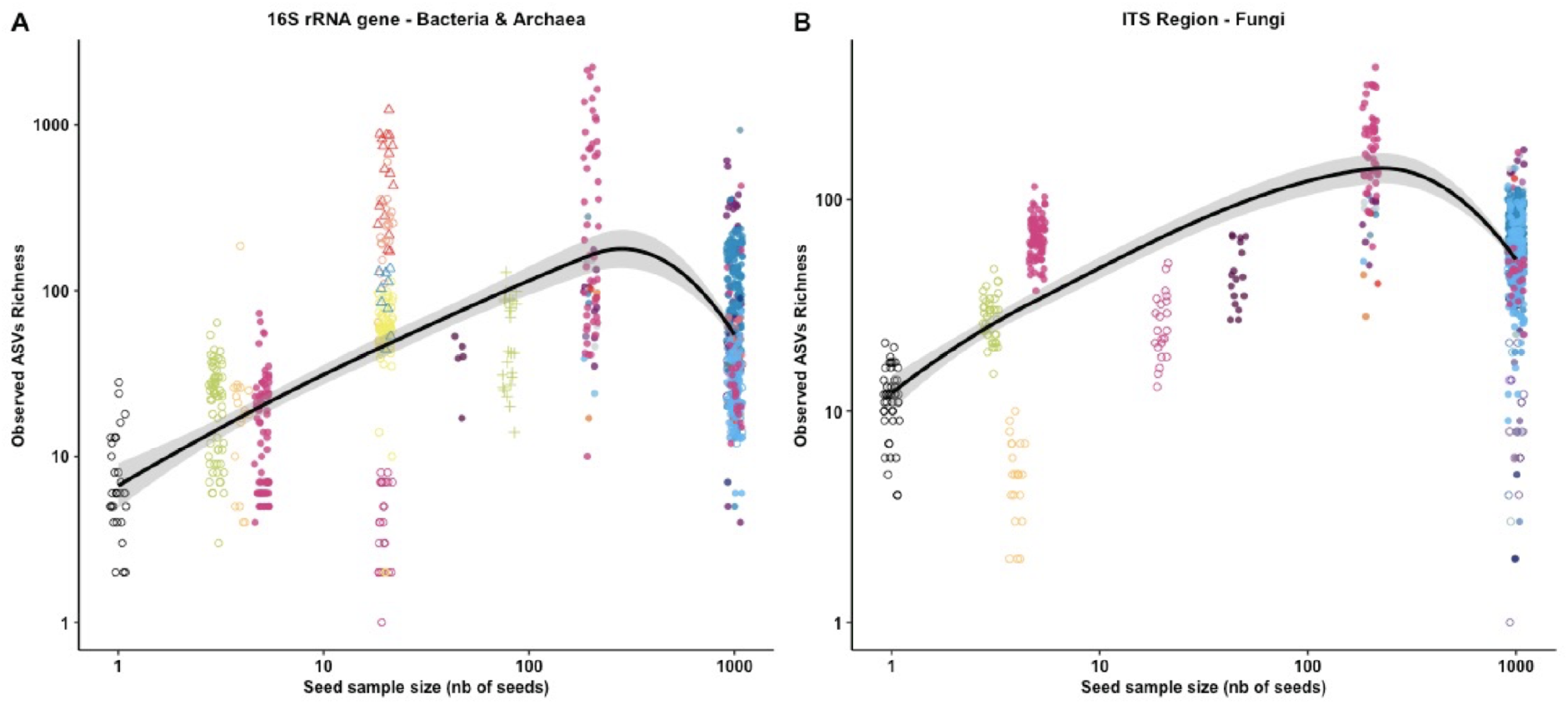
Relationship between the number of seeds included in sample and the ASV richness observed, for both 16S rRNA gene - V4 dataset (A) and the ITS1 dataset (B). The points are colored by the plant species (see legend in Figure S7).

**Figure S9:**
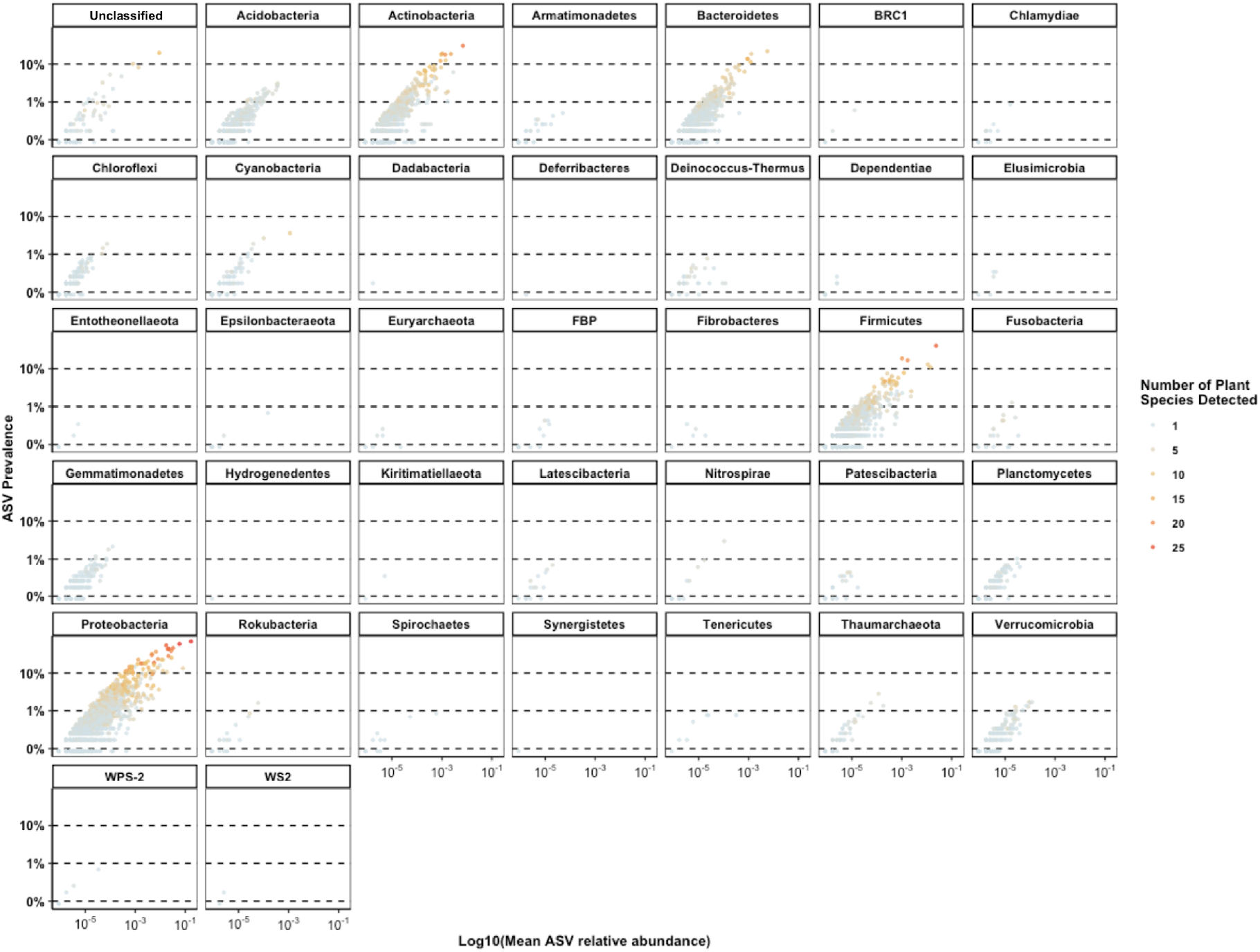
Abundance-occupancy curves for all prokaryotic phyla (Bacteria and Archaea) for the 16S rRNA gene - V4 dataset. Each point represents an ASV and the points are colored based on the number of plant species in which they were detected.

**Figure S10:**
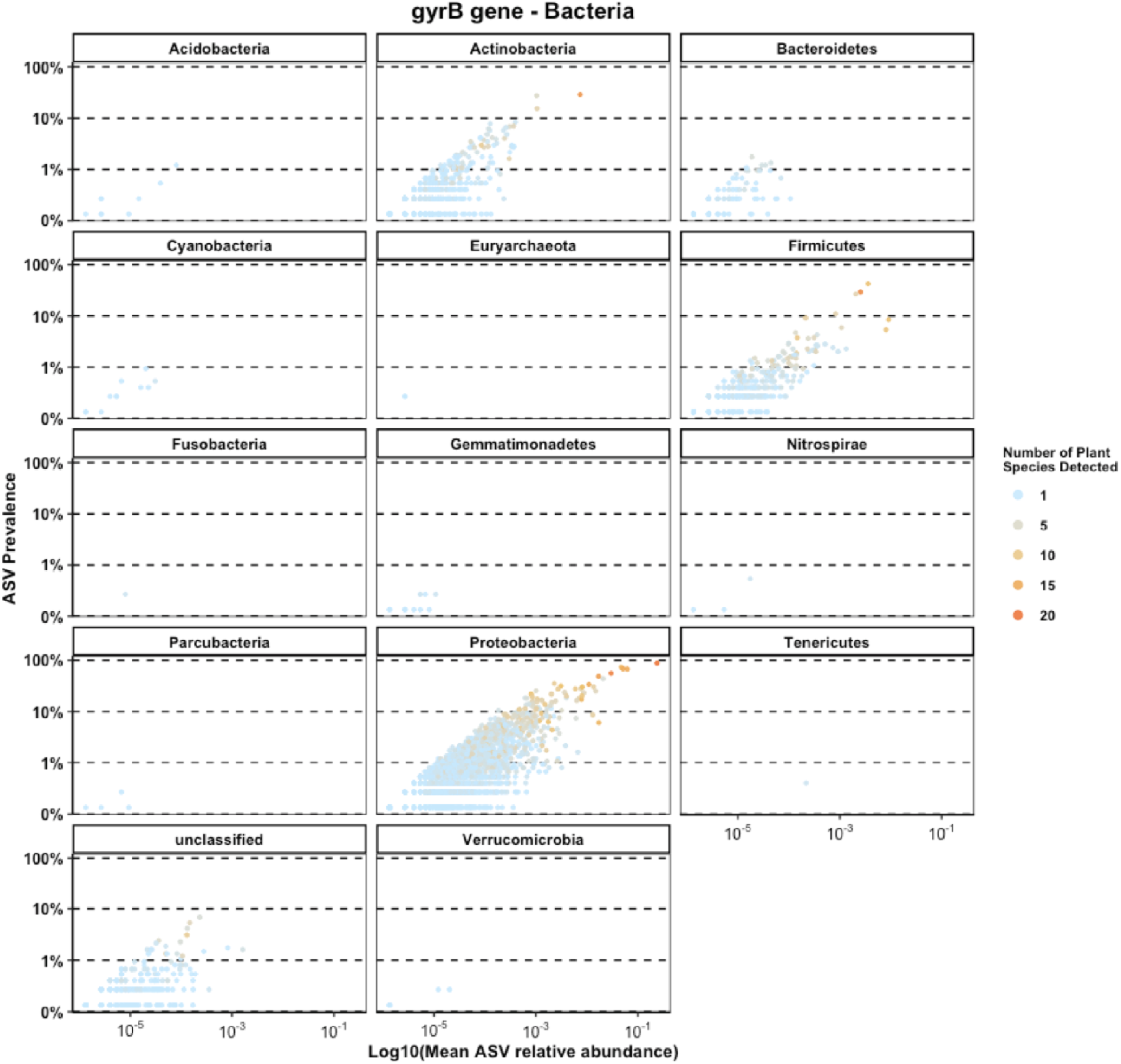
Abundance-occupancy curves for all prokaryotic phyla (Bacteria and Archaea) for the gyrB gene dataset. Each point represents an ASV and the points are colored based on the number of plant species in which they were detected.

**Figure S11:**
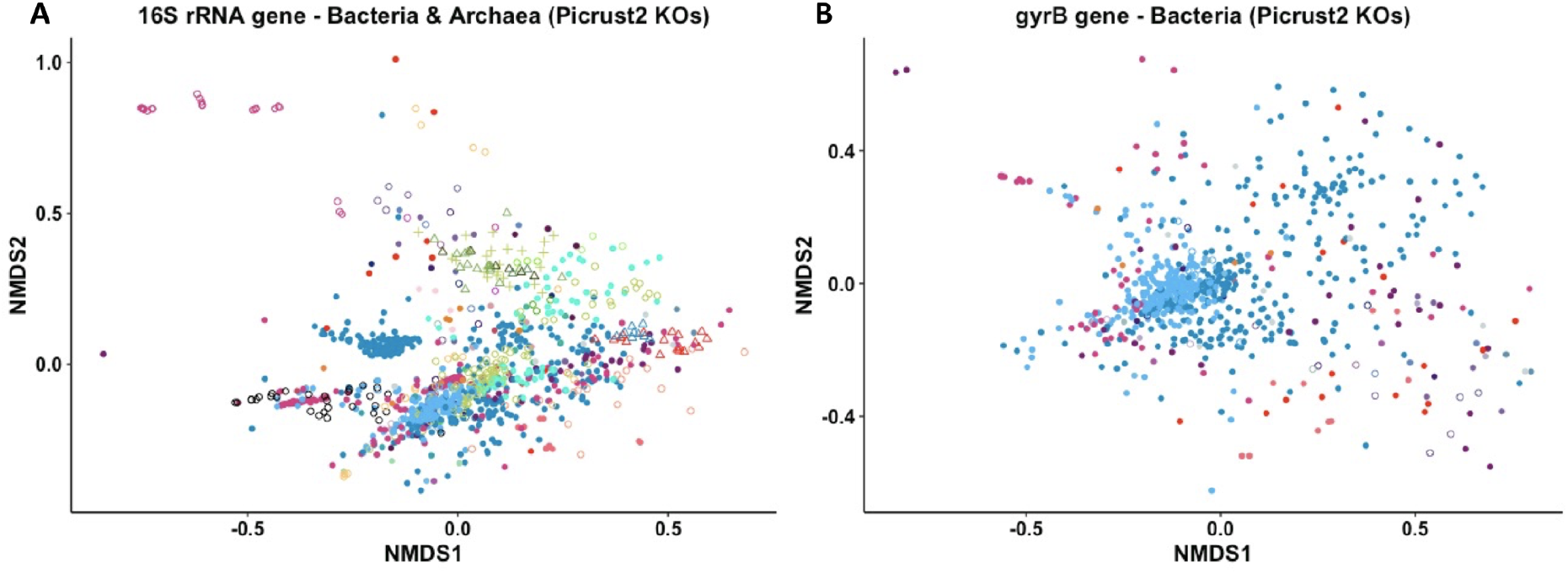
Metagenome prediction of seed microbiota based on KEGG orthologs (KOs) with Picrust2, A) for the 16S-V4 dataset and B) for the *gyrB* dataset. For colors, see legend in Figure S7

